# Retinal microglia exacerbate uveitis by functioning as local antigen-presenting cells

**DOI:** 10.1101/2024.03.23.586440

**Authors:** Shintaro Shirahama, Yoko Okunuki, May Y. Lee, Margarete M. Karg, Nasrin Refaian, Drenushe Krasniqi, Kip M. Connor, Meredith S. Gregory-Ksander, Bruce R. Ksander

## Abstract

Autoimmune uveitis is a major cause of blindness in the working-age population of developed countries. Experimental autoimmune uveitis (EAU) depends on activation of interphotoreceptor retinoid-binding protein (IRBP) specific CD4^+^ effector T cells that migrate systemically and infiltrate into the retina. Following systemic induction of retinal antigen-specific T cells, the development of EAU can be broken down into three phases: early phase when inflammatory cells begin to infiltrate the retina, amplification phase, and peak phase. Although studied extensively, the function of local antigen-presenting cells (APCs) within the retina remains unclear. Two potential types of APCs are present during uveitis, resident microglia and infiltrating CD11c^+^ dendritic cells (DCs). MHC class II (MHC II) is expressed within the retina on both CD11c^+^ DCs and microglia during the amplification phase of EAU. Therefore, we used microglia specific (P2RY12 and TMEM119) and CD11c^+^ DC specific MHC II knockout mice to study the function of APCs within the retina using the conventional and adoptive transfer methods of inducing EAU. Microglia were essential during all phases of EAU development: the early phase when microglia were MHC Il negative, and amplification and peak phases when microglia were MHC II positive. Unexpectedly, retinal infiltrating MHC Il^+^ CD11c^+^ DCs were present within the retina but their antigen-presenting function was not required for all phases of uveitis. Our data indicate microglia are the critical APCs within the retina and an important therapeutic target that can prevent and/or diminish uveitis even in the presence of circulating IRBP-specific CD4^+^ effector T cells.

## Introduction

Uveitis is one of the major causes of blindness in the working-age population (20-60 years old) and is estimated to be responsible for 10-15% of all cases of blindness in the developed world (1–3). Moreover, the loss of vision in the young working-age population is associated with a significant negative economic impact (4,5). Although autoimmune uveitis comprises a variety of clinical entities, an autoimmune response to retinal antigens is considered fundamental to its pathogenesis (6). The immunological mechanisms of autoimmune uveitis have been studied extensively in animal models of experimental autoimmune uveitis (EAU) that are induced by systemic immunization with retinal autoantigens such as interphotoreceptor retinoid-binding protein (IRBP) in mice and immunization with S-antigen (Arrestin) in rats, both EAU models require that the autoantigens are mixed with adjuvants to trigger activation of innate immunity (7–9). Following systemic induction of retinal antigen-specific T cells, the development of EAU can be broken down into three phases: the early phase, when inflammatory cells begin to infiltrate the retina, occurs 7-14 days post-immunization, followed by the amplification phase on days 14-21, culminating in the peak phase of retinitis on days 21-28.

EAU is mediated by autoantigen-specific T cells and two subsets of CD4^+^ helper T (Th) cells are known to function as effectors, interleukin (IL)-12 driven Th1 cells that release interferon gamma (IFN-γ) and IL-23 driven Th17 cells that release the potent proinflammatory cytokine IL-17 (10,11). Th17 cells have a dominant role in uveitis, as demonstrated by the ability of anti-IL-17 blocking antibodies to limit the development of EAU (12, 13). However, Th1 cells are also required to activate Th17 cells, and uveitis still develops in IL-17 knockout mice, indicating a dual role of Th1 / Th17 cells in mediating EAU (10,11). Extensive research of EAU models for many decades has revealed a great deal of information about the immune mechanisms of disease pathogenesis. However, despite these many important discoveries, there are still significant gaps in our knowledge. In particular, the function of antigen-presenting cells (APCs) within the retina that present autoantigens in the context of MHC class II (MHC II) to T cell receptors on retinal infiltrating autoantigen-specific Th1 / Th17 cells.

Over 25 years ago, Robert *et al.* discovered that the blood-retinal barrier prevented infiltration of naïve T cells but did not prevent infiltration of activated T cells (14). Moreover, retinal infiltration of activated T cells was independent of whether the T cells were specific for retinal autoantigens, as shown by the fact that activated non-retinal specific T cells (ConA activated) infiltrated into the retina at the same rate as activated S-antigen-specific T cells, at least for the first 24 hrs (14). This indicated that recognition of antigen presented by MHC II was not a requirement to gain access into the retina, only that the T cells were activated. Although T cells specific for non-retinal antigens initially infiltrate the retina, they do not persist, and the number of these T cells gradually diminishes without any evidence of retinal inflammation. By contrast, retinal autoantigen-specific T cells infiltrate the retina and are retained but do not immediately initiate retinal inflammation during the first 24-48 hrs (14). During this time, it is believed that autoantigen-specific T cells recognize their cognate antigen presented by MHC II^+^ APCs, which triggers reactivation and expansion of the antigen-specific T cells. These activated T cells produce inflammatory cytokines and chemokines, leading to the breakdown of the blood-retinal barrier and recruitment of additional inflammatory cells (T cells, monocytes, and neutrophils), resulting in retinal tissue damage (15,16). Therefore, a critical step in whether disease develops is determined, in part, by the first encounter of T cells with autoantigens presented by MHC II^+^ APCs within the retina. For this reason, identifying and determining the function of MHC II^+^ APCs in the retina is critical to understanding the pathogenesis of EAU.

There are two potential sources of MHC II^+^ APCs within the retina during the development of uveitis: activated microglia and infiltrating CD11c^+^ dendritic cells (DCs). In a normal homeostatic retina, there are few, if any, MHC II^+^ cells. Microglia only express MHC II after they are activated, and infiltrating MHC II^+^ macrophages are only present within the retina after pro-inflammatory cytokines break down the blood-retinal barrier (16, 17). Past studies of MHC II^+^ APCs within the retina during uveitis were hindered by the inability to distinguish microglia from infiltrating macrophages since, until recently, there were no cell surface markers that could distinguish microglia from infiltrating macrophages. However, new markers enriched on microglia have now been identified, such as P2RY12 (18) and TMEM119 (19), leading to new possibilities to identify and study the function of MHC II^+^ APCs during uveitis.

In our previous study, we were able to eliminate microglia in the retina *in vivo* using PLX5622, an antagonist of the colony-stimulating factor-1 receptor (CSF1R), which is highly expressed on microglia and signaling through this receptor is required for microglia survival (20–22). Using the conventional model of EAU to induce IRBP-specific CD4^+^ T cells, we demonstrated that elimination of retinal microglia by PLX5622 treatment blocked development of EAU, indicating that the initiation of EAU was dependent upon the presence of microglia (21). However, these studies were not able to specifically examine the MHC II -dependent and independent functions of microglia, as well as the APC functions of retinal infiltrating DCs. In the current study, we used the Cre-lox approach to deplete MHC II on either microglia or DCs specifically. To deplete MHC II on microglia, *P2ry12^CreER/+^*mice were crossed with homozygous loxP-flanked MHC II mice (*MHC II^fl/fl^*) to generate mice in which tamoxifen treatment induces depletion of MHC II specifically on microglia. Because several reports indicate that P2RY12 may be downregulated on activated microglia (18, 23, 24), we also crossed *MHC II^fl/fl^* mice with *Tmem119^CreERT2/+^*mice to obtain a second mouse line in which tamoxifen treatment can specifically induce MHC II depletion on microglia. A third mouse line was created to deplete MHC II specifically on DCs by crossing *MHC II^fl/fl^*mice with *CD11c^Cre^*^/0^ mice. Using these transgenic mouse lines, we studied the role of MHC II positive and negative microglia and DCs within the retina using the conventional and adoptive transfer methods of inducing EAU. Our results indicate that microglia are essential during all three phases of EAU development: in the early phase, when microglia are MHC II negative, and in the amplification and peak phases, when microglia are MHC II positive. Furthermore, our data support that MHC II^+^ microglia are the initial APCs within the retina that trigger disease. Unexpectedly, retinal infiltrating MHC II^+^ CD11c^+^ DCs were present within the retina, but their Class II antigen-presenting function was not required for all three phases of uveitis. Thus, microglia are the critical APCs within the retina and an important therapeutic target that can prevent and/or diminish uveitis even in the presence of circulating IRBP-specific CD4^+^ T cells.

## Results

### MHC II upregulation occurs within the retina on both CD11c^+^ dendritic cells and microglia during the amplification phase of EAU

MHC II functions as an antigen-presenting complex, and is critical for the function of APCs and the initiation of antigen-specific immune responses (25). While the function of MHC II^+^ CD11c^+^ DCs in the activation and expansion of CD4^+^ T cells in the spleen and lymph nodes has been studied extensively in EAU, it is still unclear which MHC II^+^ APC initiates and amplifies destructive inflammation within the retina. To begin to answer this question, conventional EAU was induced in wild-type (WT) C57BL/6J mice via immunization with the retinal antigen IRBP (26). Following systemic induction of IRBP-specific T cells, the development of uveitis can be divided into three phases: (i) early phase, (ii) amplification phase, and (iii) peak phase (27). Retinal inflammation begins at 7-14 days after immunization (early phase), gradually increases between 14-21 days (amplification phase), and peaks between 21-28 days (peak phase) (Fig. 1a). To evaluate the percentage and number of MHC II^+^ cells in the retina during EAU development, retinas were collected at day 10 (early phase) and day 20 (amplification phase) post EAU induction. At 20 days post EAU induction, there was a significant increase in the percentage of MHC II^+^ cells within the retina as compared to 10 days post EAU induction and naïve mice (*p*<0.0001, Fig. 1b and 1c). The total number of MHC II^+^ cells within the retina increased 18-fold at 20 days after EAU induction (2324.5±166.1) as compared to 10 days post EAU induction (126.0±29.6) and naïve mice (128.8±18.6) (Fig. 1b and 1c). Additional staining for CD11c (DCs and microglia) and TMEM119 (microglia-specific), revealed the MHC II^+^ cells at day 20 post EAU induction were comprised of equal numbers of CD11c^+^ TMEM119^Neg^ DCs (34.1±0.5%) and CD11c^+^ TMEM119^+^ retinal microglia (30.3±0.8%) (Fig. 1d and 1e). These results demonstrate that CD11c^+^ DCs and retinal microglia contribute equally to the significant increase in MHC II expression observed in the retina during the amplification phase of EAU.

**Figure 1.**
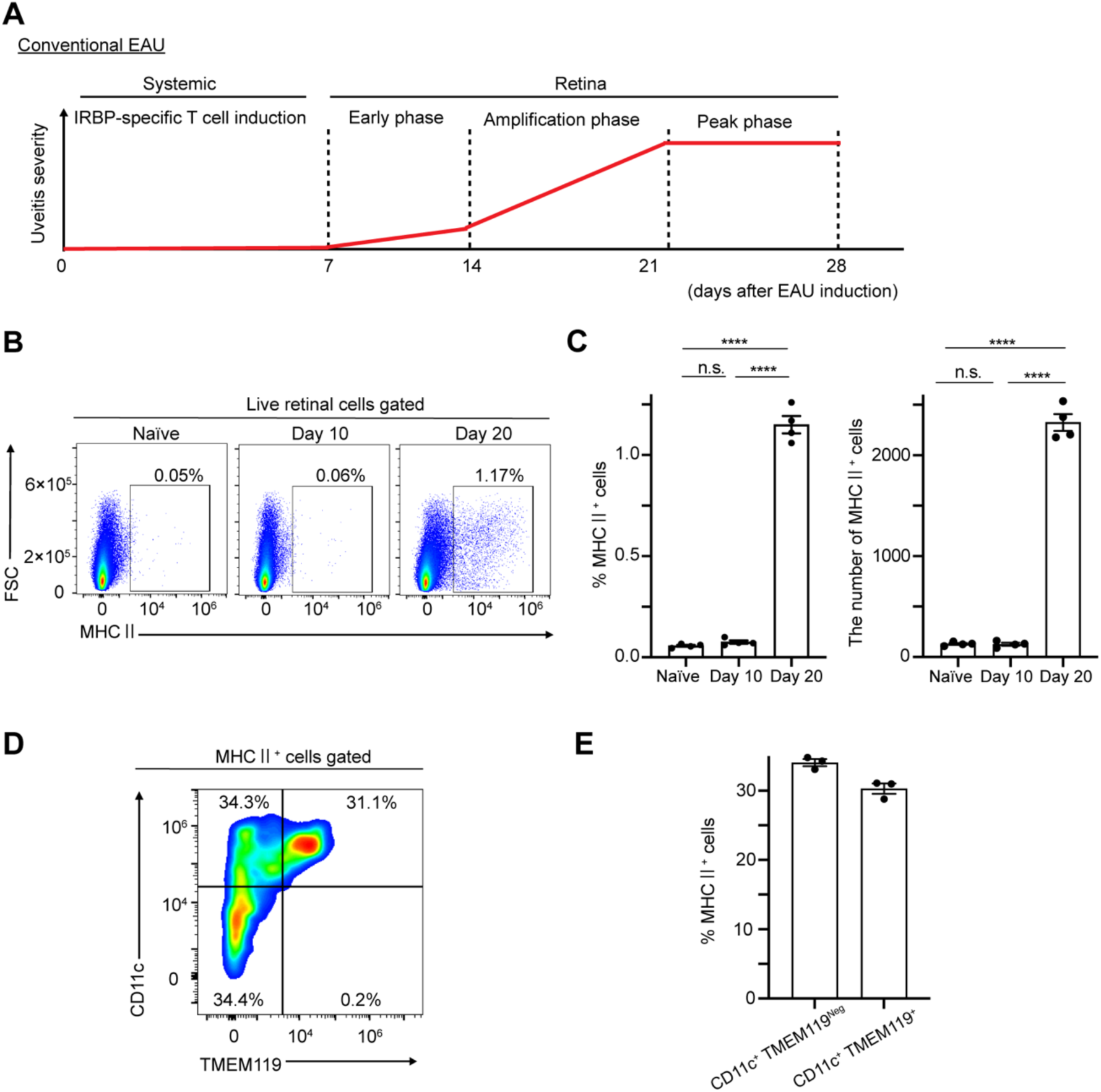
During the amplification stage of EAU, MHC II^+^ retinal cells comprise of both infiltrating dendritic cells and resident microglia. C57BL/6J mice were immunized with IRBP and at 10- and 20-days post immunization, retinas were collected and single-cell suspensions, excluding dead cells (DAPI^+^) were analyzed by flow cytometry. (A) Schematic illustrates the three stages of conventional experimental autoimmune uveitis (EAU); Early phase, Amplification phase, and Peak phase. (B) Representative flow cytometry plots showing MHC II expression in the retina from naïve, day 10 post immunization, and day 20 post immunization. (C) Quantitative analysis of the MHC II population (naïve, n=4; day 10, n=4; day 20, n=4). (D) Representative flow cytometry plot showing MHC II expression on CD11c^+^ TMEM119^+^ and CD11c^+^ TMEM119^Neg^ retinal cells at 20 days post immunization (E) Quantitative analysis of the MHC II^+^ population (n=3). All data are presented as mean ± SEM. All statistical analyses were performed using a one-way analysis of variance with Tukey’s multiple comparison test. \*\*\*\**p*<0.0001.

To look more closely at retinal microglia during the course of EAU development, flow cytometry was performed with two microglia-specific markers (P2RY12 and TMEM119) to quantitate the number of retinal microglia, as well as the level of MHC II expression. Conventional EAU was induced in WT C57BL/6J mice, and flow cytometry was performed on naïve retinas and retinas isolated on day 10 and day 20 post EAU induction. To identify retinal microglia, cells were first gated based on CD45 and CD11b expression, where microglia are expected to be CD45^low^ CD11b^+^ (28). Two additional microglia-specific markers (P2RY12 and TMEM119) were then used to define further the retinal microglia population (Supplementary Fig. 1a and 1b). Both P2RY12 and TMEM119 staining reveal a significant increase in the total number of retinal microglia at 20 days post EAU induction (1172.7±121.7 and 809.5±90.4, respectively) as compared to 10 days post EAU induction (643.0±176.4 and 588.3±36.9, respectively) and naïve retinas (628.7±141.6 and 458.1±118.3, respectively) (Fig. 2b and 2e). In addition, at 20 days post EAU induction, MHC II expression was significantly increased in these P2RY12^+^ and TMEM119^+^ microglia, with 55.9±5.6% of P2RY12+ cells and 61.5±5.8% of TMEM119^+^ cells expressing MHC II at 20 days post EAU induction, as compared to only 1.8±0.5% of P2RY12^+^ and 2.9±0.6% of TMEM119^+^ cells expressing MHC II at 10 days post EAU induction and 1.9±0.5% of P2RY12^+^ and 1.8±0.3% of TMEM119^+^ cells expressing MHC II in naïve (Fig. 2a, 2c, 2e, and 2f). As a result, there was a 57-fold increase in the number of MHC II^+^ P2RY12^+^ retinal microglia at 20 days post EAU induction (651.3±40.8) when compared to 10 days post EAU induction (11.5±2.6) and naïve (11.5±2.4) (Fig. 2c). Similar results were observed using TMEM119, with a 29-fold increase in the number of MHC II^+^ TMEM119^+^ retinal microglia at 20 days post EAU induction (494.8±28.5) when compared to 10 days post EAU induction (17.0±4.4) and 32-fold increase when compared to naïve (15.5±3.9) (Fig. 2f). In contrast, there was no significant difference in the total number of P2RY12^+^ and TMEM119^+^ retinal microglia nor the percentage of MHC II ^+^retinal microglia between naïve and 10 days post EAU induction. These results demonstrate that retinal microglia remain MHC II^Neg^ during the early phase of EAU, followed by a significant increase in the number of microglia and MHC II^+^ expression during the amplification phase of EAU.

**Figure 2.**
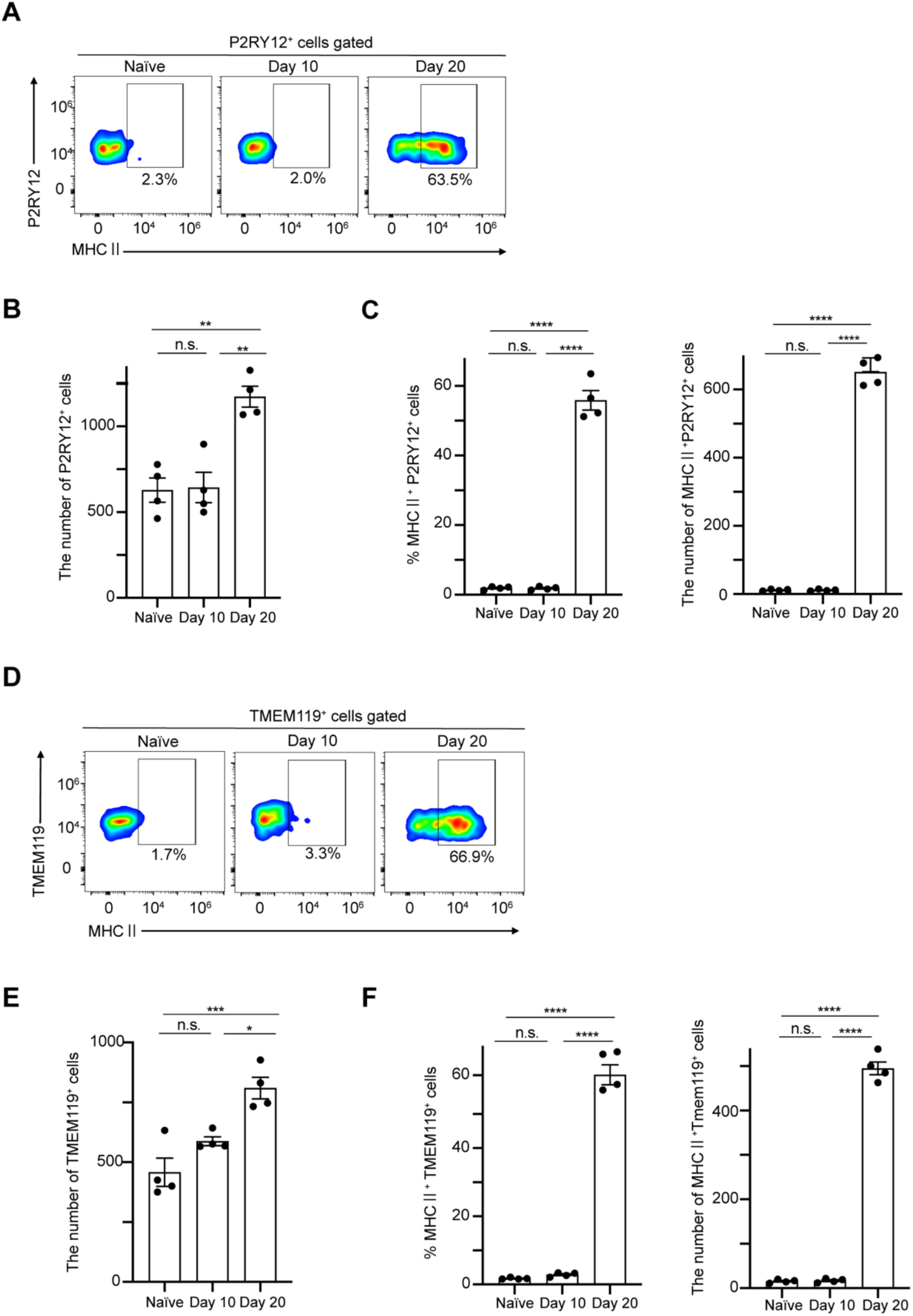
The number of retinal microglia and expression of MHC II increases significantly during the amplification stage of EAU. C57BL/6J mice were immunized with IRBP and at 10- and 20-days post immunization, retinas were collected and single-cell suspensions, excluding dead cells (DAPI^+^) were analyzed by flow cytometry. (A) Representative flow cytometry plots showing MHC II expression on P2RY12^+^ cells in the retina from naïve, day 10 post immunization, and day 20 post immunization. (B) Quantitative analysis of the P2RY12^+^ population (naïve, n=4; day 10, n=4; day 20, n=4). (C) Quantitative analysis of the MHC II^+^ P2RY12^+^ population (naïve, n=4; day 10, n=4; day 20, n=4). (D) Representative flow cytometry plots showing MHC II expression on TMEM119^+^ cells in the retina from naïve, day 10 post immunization, and day 20 post immunization. (E) Quantitative analysis of the TMEM119^+^ population (naïve, n=4; day 10, n=4; day 20, n=4). (F) Quantitative analysis of the MHC II^+^ TMEM119^+^ population (naïve, n=4; day 10, n=4; day 20, n=4). All data are presented as mean ± SEM. All statistical analyses were performed using a one-way analysis of variance with Tukey’s multiple comparison test. \**p*<0.05; \*\**p*<0.01; \*\*\*\**p*<0.001; \*\*\*\**p*<0.0001.

### MHCII^+^ retinal microglia are required for the amplification and peak phases, but not for the early phase of EAU

To elucidate the function of MHC II^+^ DCs and MHC II^+^ retinal microglia in the different phases of EAU, we used the Cre-lox approach to specifically deplete MHC II on microglia and DCs. To deplete MHC II on microglia, we crossed *P2ry12^CreER/+^* mice with homozygous loxP-flanked MHC II mice (*MHC II^fl/fl^*) to generate mice in which tamoxifen treatment induces depletion of MHC II specifically on retinal microglia. Because several reports indicate that P2RY12 may be downregulated on activated microglia (18, 23, 24), we also crossed *MHC II ^fl/fl^* mice with *Tmem119^CreERT2/+^*mice to obtain a second mouse line in which we can specifically induce MHC II depletion on microglia. A third mouse line was created to deplete MHC II on DCs by crossing *MHC II ^fl/fl^* mice with *CD11c^Cr/0^*mice.

To confirm knockdown of MHC II on retinal microglia of *MHC II^fl/fl^ P2ry12^CreER/+^* mice and *MHC II^fl/fl^ Tmem119^CreERT2/+^*mice, we injected tamoxifen intraperitoneally for five consecutive days, followed by induction of conventional EAU (Supplementary Fig. 2a). At 21 days post EAU induction, retinas were isolated and microglia-specific MHC II expression was assessed by flow cytometry. In Experiment 1, the percentage of P2RY12^+^ microglia expressing MHC II in tamoxifen-treated *MHC II^fl/fl^ P2ry12^CreER/+^*mice was reduced by approximately 74% as compared to tamoxifen-treated control mice *MHC II^fl/fl^ P2ry12^+/+^* mice (no Cre) (12.4±2.4% vs 46.9±3.4 %) (Supplementary Fig. 2b).

Similarly, in Experiment 2, the percentage of TMEM119^+^ microglia expressing MHC II in tamoxifen-treated *MHC II^fl/fl^ Tmem119^CreERT2/+^* mice was reduced by approximately 73% as compared to tamoxifen-treated control mice *MHC II^fl/fl^ Tmem119^+/+^*mice (no Cre) (13.4±2.6% vs 49.4±5.5 %) (Supplementary Fig. 2c). To confirm knockdown of MHC II on CD11c^+^ DCs in *MHC II^fl/fl^ CD11c^Cre/0^* mice, draining lymph nodes (LNs) and spleen (SP) were isolated at 21 days post EAU and MHC II expression was assessed by flow cytometry (Supplementary Fig. 3a). DCs were first identified as CD45^+^ CD11c^+^ and at 21 days post EAU induction the percent of CD45^+^ CD11c^+^ DCs expressing MHC II in *MHC II^fl/fl^ CD11c^Cre/0^* mice was reduced by approximately 90% compared to that in *MHC II^fl/fl^ CD11c^+/0^* mice (7.3±1.7 % vs 75.9±1.9 %), indicating a successful depletion of MHC II expression on CD11c^+^ DCs (Supplementary Fig. 3b, 3c, and 3d).

To assess the effects of MHC II depletion on EAU development, conventional EAU was induced in tamoxifen-treated *MHC II^fl/fl^ P2ry12^+/+^* (control mice) and *MHC II^fl/fl^ P2ry12^CreER/+^*mice (Fig. 3a). Clinical assessment of EAU pathology by fundus examination was performed at 7,14, 21-, and 28-days post EAU induction (Fig. 3b) and histological assessment via H&E staining was performed at 21 days post EAU induction (Fig. 3c). In the absence of MHC II expression on P2RY12^+^ microglia, fundus examination revealed a significant decrease in retinal inflammation at day 21 (amplification phase) and day 28 (peak phase) post EAU induction as compared to tamoxifen treated control mice (*MHC II^fl/fl^ P2ry12^+/+^*). However, no significant difference in retinal inflammation was observed during the early phase (day 7-14) (Fig. 3b). Histological evaluation on day 21 further confirmed that P2RY12-specific depletion of MHC II significantly suppressed retinal inflammation during the amplification phase of EAU (Fig. 3c).

**Figure 3.**
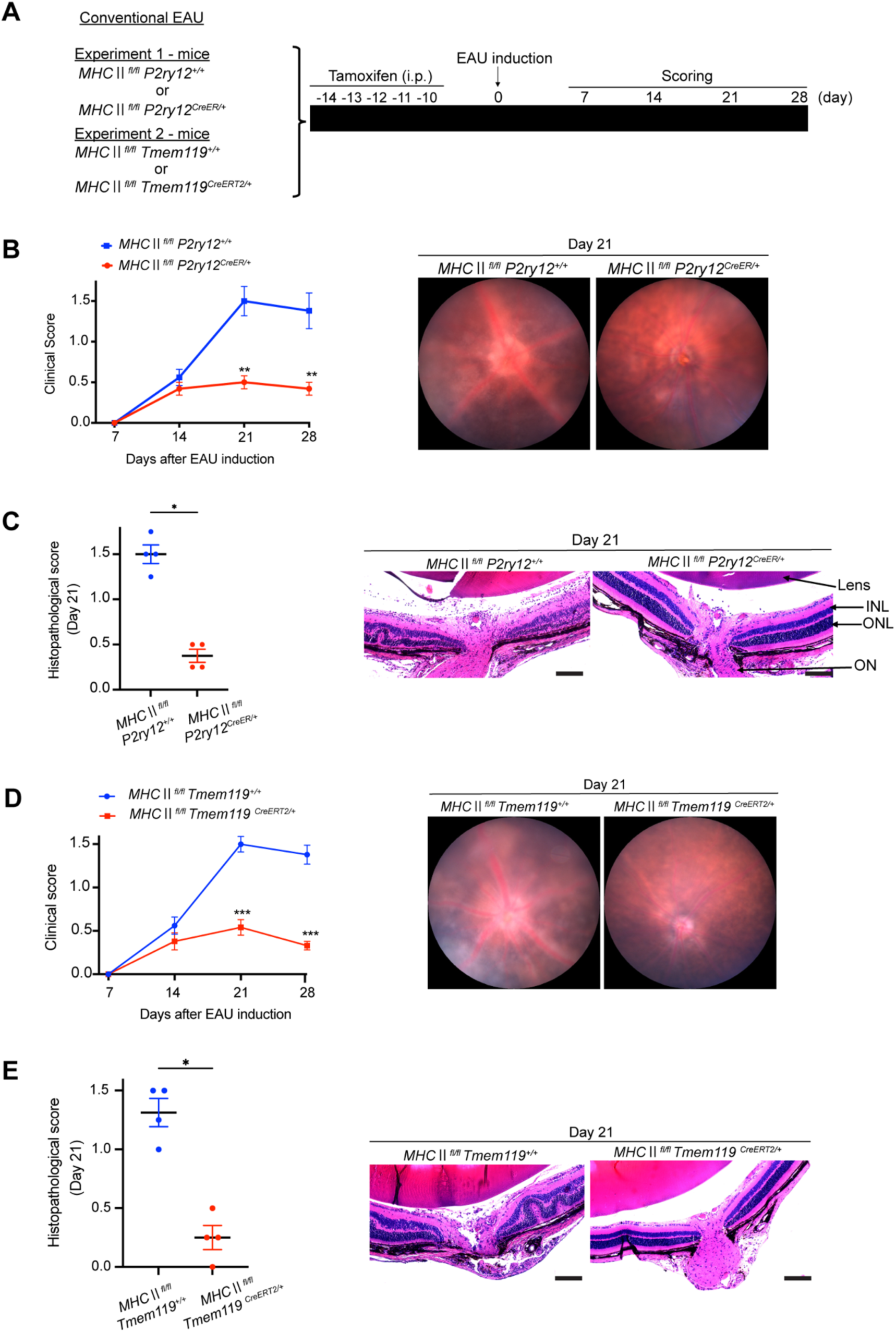
MHC II^+^ expression on retinal microglia is required for the amplification of retinal inflammation during EAU. (A) Schematic time course of conventional experimental autoimmune uveitis (EAU) experiment in which Tamoxifen was administration was initiated 14 days prior to immunization with IRBP in order to specifically knock out MHC II on microglia, using *MHC II^fl/fl^ P2ry12^CreER/+^* and *MHC II^fl/fl^ Tmem119^CreER/+^* mice. (B) The time course of EAU clinical scores for *MHC II^fl/fl^ P2ry12^+/+^* (n=4) and *MHC II^fl/fl^ P2ry12^CreER/+^*(n=6) mice and representative fundus images. (C) Histopathological EAU score on day 21 post EAU induction and representative histopathological images (H&E staining) (n=4 per group). Scale bar: 100 μm. (D) The time course of EAU clinical scores for *MHC II^fl/fl^ Tmem119^+/+^* (n=4) and *MHC II^fl/fl^ Tmem119^CreER/+^*(n=6) mice and representative fundus images. (E) Histopathological EAU score for *MHC II^fl/fl^ Tmem119^+/+^* and *MHC II^fl/fl^ Tmem119^CreER/+^* mice on day 21 post EAU induction and representative histopathological images (H&E staining) (n=5 per group). Scale bar: 100 μm. All data are presented as mean ± SEM. All statistical analyses were performed using the Mann-Whitney U test. \**p*<0.05; \*\**p*<0.01; \*\*\**p*<0.001.

Similar results were obtained following induction of conventional EAU in tamoxifen-treated *MHC II^fl/fl^ Tmem119^+/+^* (control mice) and *MHC II^fl/fl^ Tmem119^CreER/+^* mice (Fig. 3a). In the absence of MHC II expression on TMEM119^+^ microglia, fundus examination revealed a significant decrease in retinal inflammation at day 21 and day 28 post EAU induction (Fig. 3d). However, no significant difference in retinal inflammation was observed during the early phase (day 7-14). Histological evaluation on day 21 further confirmed that TMEM119-specific depletion of MHC II significantly suppressed retinal inflammation during the amplification phase of EAU (Fig. 3e). Together, these data demonstrate that MHC II ^+^ retinal microglia mediate the amplification and peak phases of EAU.

### Increased MHC II expression on retinal microglia correlates with increased EAU severity

To further examine the relationship between MHC II expression on retinal microglia and the severity of uveitis, conventional EAU was induced in both tamoxifen-treated *MHC II^fl/fl^ P2ry12^CreER/+^*mice and *MHC II^fl/fl^ Tmem119^CreERT2/+^* mice (Fig. 4a). *MHC II^fl/fl^ P2ry12^+/+^* and *MHC II^fl/fl^ Tmem119^+/+^* mice served as MHC II positive controls. At 21 days post EAU induction, fundus examination was performed to assess EAU severity. At the same time, the retinas were isolated and microglia-specific MHC II expression was assessed by flow cytometry (Fig. 4b-4e). A correlation analysis between clinical score and the percentage of MHC II^+^ microglia in the retina revealed a strong positive correlation (r=0.9696, *p*<0.0001) between the percentage of MHCII^+^ P2RY12^+^ cells in the retina and the clinical score (Fig.4b). Similarly, a strong positive correlation (r=0.9482, *p*<0.0001) was also observed between the percentage of MHCII^+^ TMEM119^+^ cells in the retina and the clinical score (Fig.4d). These results further confirm the importance of MHC II ^+^ retinal microglia in the amplification of retinal inflammation in EAU, with increased expression of MHC II on retinal microglia correlating with increased severity of EAU.

**Figure 4.**
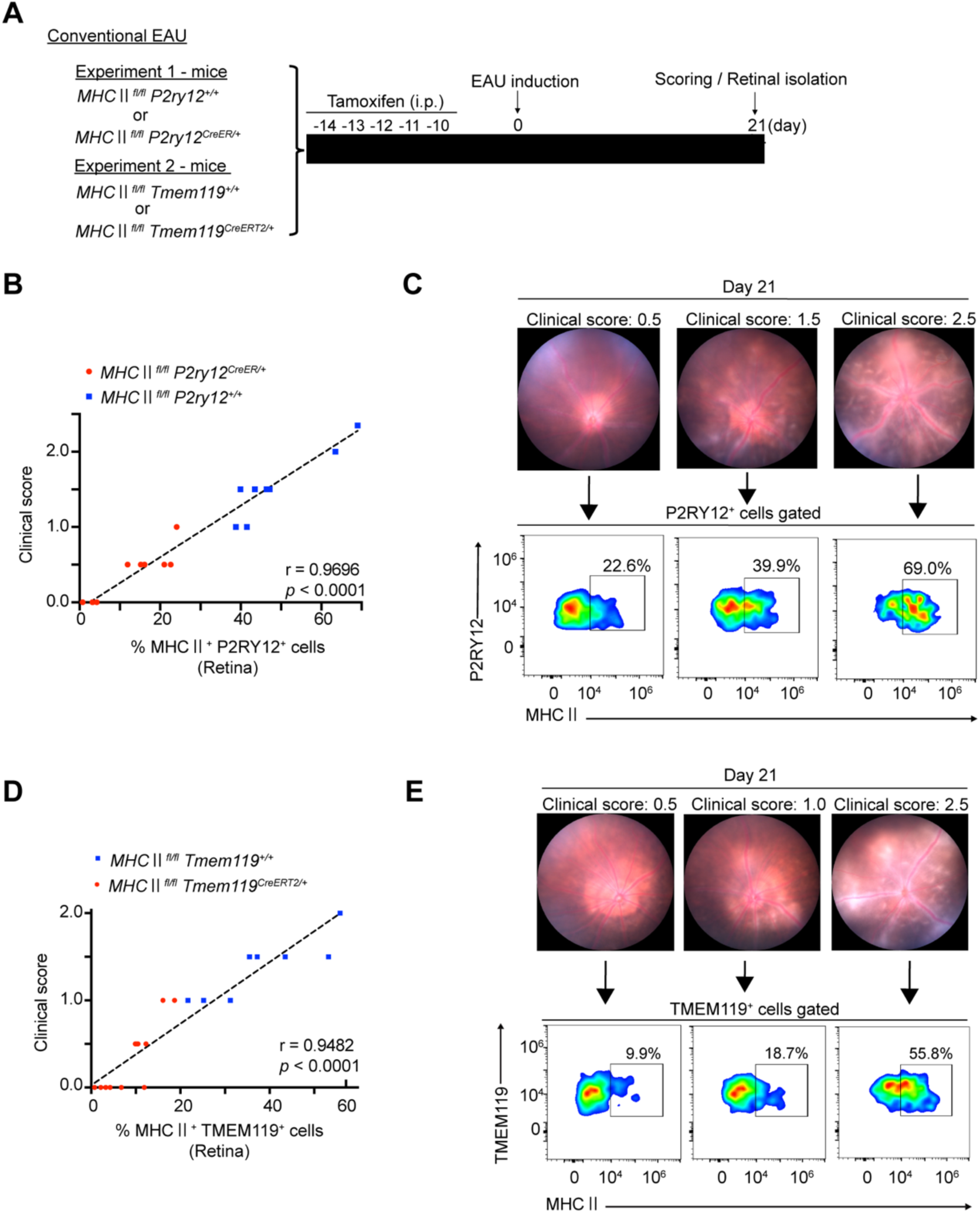
Increased expression of MHC II on retinal microglia correlates with increased severity of EAU. (A) Schematic time course of conventional EAU experiment in which Tamoxifen administration was initiated 14 days before immunization with IRBP to specifically knock out MHC II on microglia, using *MHC II^fl/fl^, P2ry12^CreER/+^* and *MHC II^fl/fl^ Tmem119^CreER/+^* mice. The clinical score was evaluated on day 21 post IRBP immunization, and the retina was collected for flow cytometry. (B) Spearman’s rank correlation coefficient was calculated to examine the relationship between clinical score and percentage of MHCⅡ^+^, P2RY12^+^ cells in the retina at day 21 post-IRBP immunization (*MHC II^fl/fl^ P2ry12^+/+^*, n=8; *MHC II^fl/fl^ P2ry12^CreER/+^*, n=10) (C) Representative fundus images (Clinical score 0.5, 1.5, and 2.5) and corresponding flow cytometry plots showing MHC II expression on P2RY12^+^ retinal cells. (D) Spearman’s rank correlation coefficient was calculated to examine the relationship between clinical score and percentage MHC II^+^ TMEM119^+^ cells in the retina at day 21 post-IRBP immunization (*MHC II^fl/fl^ Tmem119^+/+^*, n=8; *MHC II^fl/fl^ Tmem119^CreERT2/+^*, n=12). (E) Representative fundus images (Clinical score 0.5, 1.5, and 2.5) and corresponding flow cytometry plots showing MHC II expression on TMEM119^+^ retinal cells. A correlation analysis was performed using the Spearman rank correlation analysis.

### MHC II ^+^ DCs are required for induction of EAU

MHC II ^+^ DCs are thought to be the primary APC responsible for the induction of CD4^+^ T cells that mediate EAU (9, 29) and inhibition of DC maturation, which prevents the upregulation of MHC II, attenuates EAU by preventing the generation of antigen-specific Th1 and Th17 cells (30, 31). We further confirmed the essential role of MHC II ^+^ DCs in the pathogenesis of EAU by inducing conventional EAU in *MHC II^fl/fl^ CD11c^+/0^* (control mice) and *MHC II^fl/fl^ CD11c^Cre/0^*mice (Fig. 5a). Clinical assessment revealed an almost complete absence of retinal inflammation at day 14, day 21, and day 28 post EAU induction in mice lacking MHC II^+^ DCs (Fig. 5b). Histological evaluation on day 21 EAU induction further confirmed that depletion of MHC II on CD11c^+^ DCs almost completely suppressed the development of retinal inflammation (Fig. 5c). The failure to induce EAU in *MHC II^fl/fl^ CD11c^Cre/0^*mice coincided with systemic immune response reduction as demonstrated by a significant reduction in the size of draining lymph nodes (LN) and spleen (SP) when compared to control mice (*MHC II^fl/fl^ CD11c^+/0^*) (Supplementary Fig. 4a). To further clarify whether this systemic immune response reduction depends on antigen specificity, we performed an ear swelling test, one of the delayed-type hypersensitivity assays, by intradermally injecting IRBP into the ear. IRBP-immunized *MHC II^fl/fl^ CD11c^Cre/0^* mice failed to swell the ear as compared to *MHC II^fl/fl^ CD11c^+/0^* mice (Supplementary Fig. 4b). Moreover, only LN and SP cultures prepared from IRBP-immunized *MHC II^fl/fl^ CD11c^+/0^* mice showed significant antigen-specific, IRBP-induced cell proliferation when compared to cultures prepared from IRBP-immunized *MHC II^fl/fl^ CD11c^Cre/0^*mice and naïve WT mice (Supplementary Fig. 4c). Similar results were observed in co-cultures containing purified CD4^+^ T cells (from LNs) (Supplementary Fig. 5a-5c) and purified CD11c^+^ DCs (from SP) (Supplementary Fig. 5d-5f). Purified MHC II^+^ DCs from IRBP-immunized *MHC II^fl/fl^ CD11c^+/0^* control mice were able to induce the proliferation of CD4^+^ T cells purified from both *MHC II^fl/fl^ CD11c^+/0^* and *MHC II^fl/fl^ CD11c^Cre/0^* mice, while there was a significant reduction in both the percentage and number of proliferating CD4^+^ T cells co-cultured with MHC II^Neg^ DCs purified from IRBP-immunized *MHC II^fl/fl^ CD11Cre^+/0^* mice (Supplementary Fig. 4d and 4e). Taken together, these data confirm that the MHC II ^+^ DCs serve as the primary APC responsible for the systemic activation and expansion of IRBP-specific CD4^+^ T cells in the conventional model of EAU.

**Figure 5.**
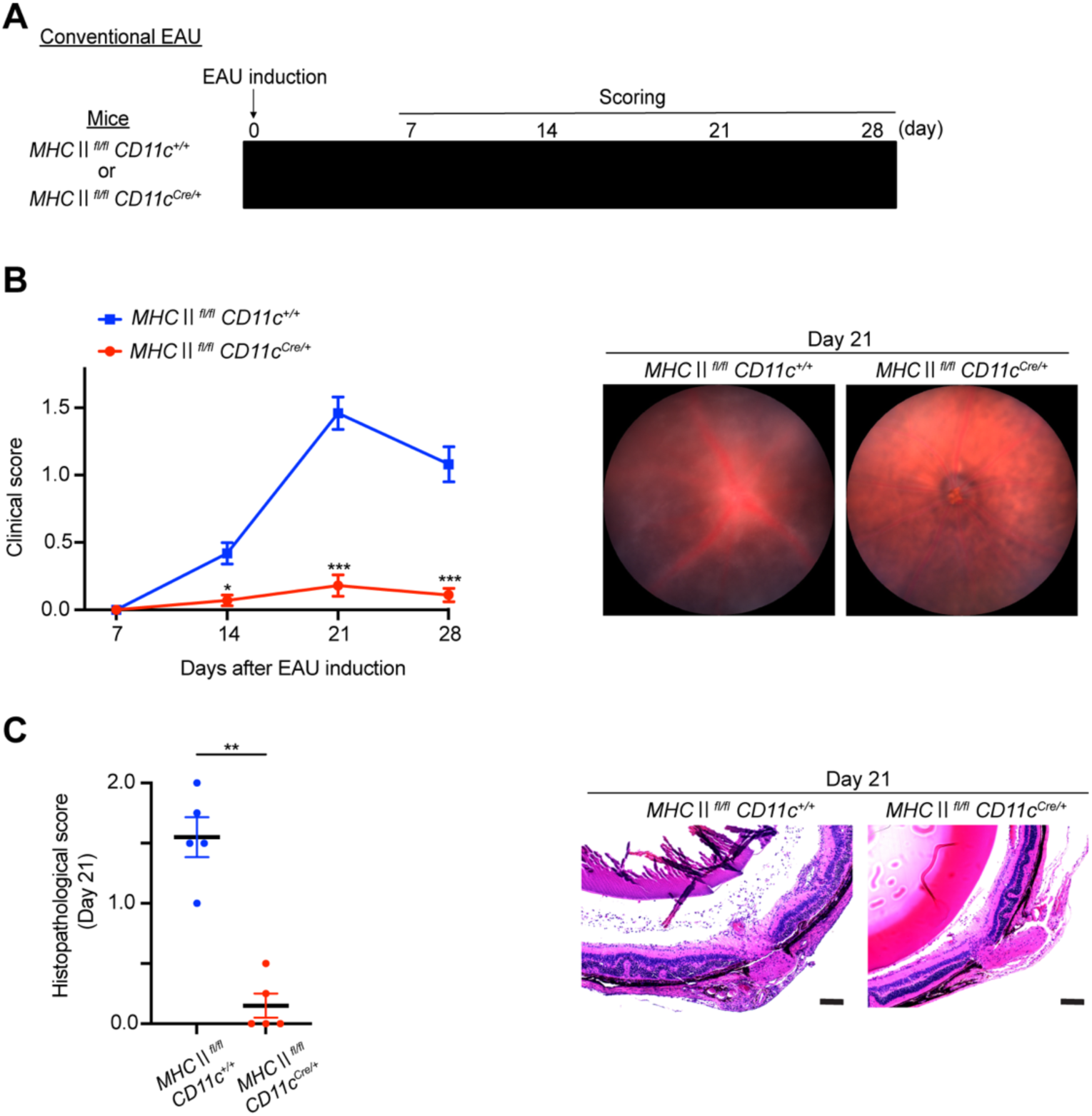
MHC II^+^ DCs are required for induction of conventional EAU. (A) Schematic time course of conventional EAU experiment in which *MHC II^fl/fl^ CD11c^+/0^* or *MHC II^fl/fl^ CD11c^Cre/0^* mice were immunized with IRBP. (B) The time course of EAU clinical scores for *MHC II^fl/fl^ CD11c^+/0^* (n=6) or *MHC II^fl/fl^ CD11c^Cre/0^* (n=7) mice and representative fundus images at day 21 post IRBP immunization. (C) Histopathological EAU score on day 21 post IRBP immunization for *MHC II^fl/fl^ CD11c^+/0^* (n=5) and *MHC II^fl/fl^ CD11c^Cre/0^* (n=5) mice and representative histopathological images (H&E staining). Scale bar: 100 μm. All data are presented as mean ± SEM. All statistical analyses were performed using the Mann-Whitney U test. \*\**p*<0.01.

### MHC II ^+^ microglia, but not MHC II ^+^ DCs, are required for the amplification and peak inflammation within the retina

In the previous experiments, uveitis did not fully develop in the absence of MHC II^+^ microglia, even though MHC II^+^ infiltrating DCs were present, indicating the local APC function of infiltrating MHC II^+^ DC is insufficient to cause disease. To further confirm this result, we wanted to eliminate MHC II on retinal infiltrating DCs and demonstrate that this has no effect on the development of uveitis. However, this experiment is only possible using the adoptive transfer model of EAU in which activated CD4^+^ T cells are generated in IRBP-immunized WT donor mice and then adoptively transferred into recipient mice that contain MHC II^Neg^ CD11c^+^ DCs (25). Like the conventional model of EAU, following adoptive transfer of IRBP-specific T cells, the development of EAU in the retina can be divided into three phases: (i) early phase, (ii) amplification phase, and (iii) peak phase (26). However, each phase occurs one week earlier in the adoptive transfer model, with retinal inflammation beginning at 0-7 days after adoptive transfer (early phase), gradually increasing between 7-14 days (amplification phase), and peaking between 14-21 days (peak phase) (Fig. 6a). Most importantly though, because EAU is induced by transferring antigen-specific lymphocytes from IRBP-immunized donor mice into recipient mice, the adoptive transfer model of EAU can be used to bypass the requirement of MHC II ^+^ CD11c^+^ DCs in recipient mice to generate the effector T cells. To illustrate this, we first collected donor cells from *MHC II ^fl/fl^ CD11c^+/0^* control mice (MHC II^+^ DCs) and *MHC II^fl/fl^ CD11c^Cre/0^*mice (MHC II^Neg^ DCs) at 14 days post EAU induction, expanded the cells for 3 days *in vitro*, and adoptively transferred them into WT recipient mice (Fig. 6b). The development of EAU was monitored via fundus examination, revealing successful induction of EAU in WT mice receiving donor cells from *MHC II^fl/fl^ CD11c^+/0^* mice (MHC II ^+^ DCs) and an almost complete failure to develop EAU in WT recipient mice receiving donor cells from *MHC II^fl/fl^ CD11c^Cre/0^* mice (MHC II^Neg^ DCs) mice (Fig. 6c). Histological EAU evaluation (day 13) further confirmed these results (Fig. 6d) and together demonstrated the requirement of MHC II ^+^ DCs in the donor mice.

**Figure 6.**
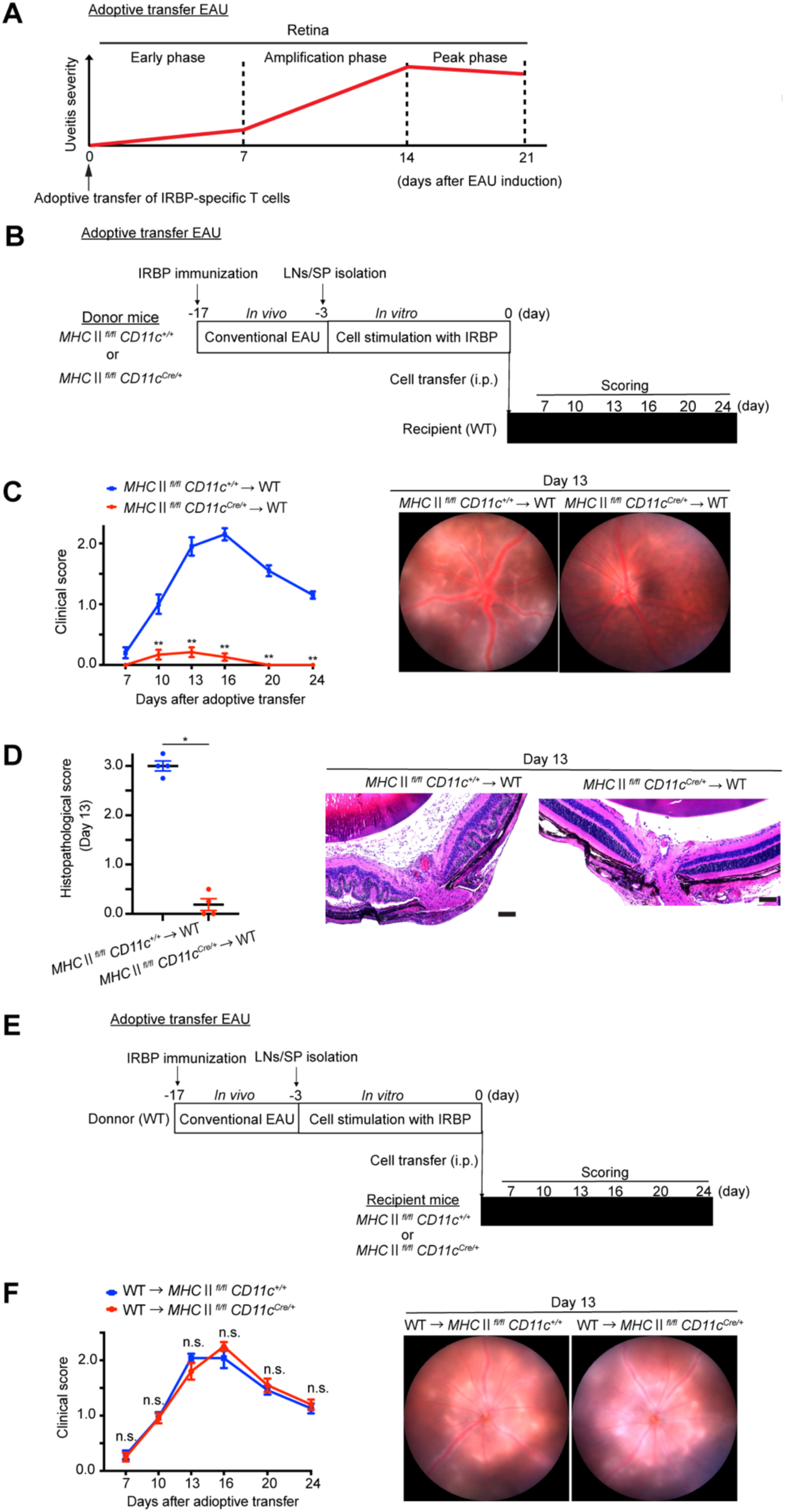
MHC II expression on CD11c^+^ DCs in recipient mice is not required to induce adoptive transfer EAU. (A) Schematic illustrates the three stages of Adoptive transfer EAU; Induction phase, Amplification phase, and Peak phase. (B) Schematic time course of adoptive transfer EAU experiment in which donor cells were isolated from *MHC II^fl/fl^ CD11c^+/0^* or *MHC II^fl/fl^ CD11c^Cre/0^*at 14 days post immunization with IRBP, expanded in vitro, and administered to C57BL/6J WT recipient mice. (C) The time course of EAU clinical scores in WT recipient mice that received donor cells from *MHC II^fl/fl^ CD11c^+/0^* (n=5) or *MHC II^fl/fl^ CD11c^Cre/0^*(n=6) mice and representative fundus images (day 13 post adoptive transfer). (D) Histopathological EAU score on day 13 post adoptive transfer and representative histopathological images (H&E staining) (n=4 per group). Scale bar: 100 μm. (E) Schematic time course of adoptive transfer EAU experiment in which donor cells were isolated from WT mice at 14 days post immunization with IRBP, expanded in vitro, and administered to *MHC II^fl/fl^ CD11c^+/0^* or *MHC II^fl/fl^ CD11c^Cre/0^* recipient mice. (F) The time course of EAU clinical scores in *MHC II^fl/fl^ CD11c^+/0^* (n=6) or *MHC II^fl/fl^ CD11c^Cre/0^* (n=5) recipient mice that received donor cells from WT mice and representative fundus images (day 13 post adoptive transfer). All data are presented as mean ± SEM. All statistical analyses were performed using the Mann-Whitney U test. \**p*<0.05; \*\**p*<0.01.

In all recipient mice, when donor cells were collected from WT mice at 14 days post EAU induction, expanded *in vitro*, and adoptively transferred into *MHC II^fl/fl^ CD11c^+/0^* control mice (MHC II^+^ DCs) and *MHC II^fl/fl^ CD11c^Cre/0^* (MHC II^Neg^ DCs) experimental mice (Fig. 6e), clinical assessment of EAU pathology by fundus examination revealed a robust EAU response irrespective of MHC II expression on recipient CD11c^+^ DCs (Fig. 6f). There was no significant reduction in uveitis when MHC II was absent on retinal infiltrating DCs, confirming those cells lack the APC function to amplify uveitis within the retina.

To also confirm the essential role of MHC II^+^ microglia in the amplification and peak phase of EAU, we transferred WT-derived activated CD4^+^ T cells into two kinds of tamoxifen-treated microglia-specific MHC II knockout mice (*MHC II^fl/fl^ P2ry12^CreER/+^* mice or *MHC II^fl/fl^ Tmem119^CreERT2/+^* mice) (Fig. 7a). Clinical assessment of EAU pathology by fundus examination was performed at 7, 10, 13, 16, 20, and 24 days post adoptive transfer and revealed a significant suppression of EAU in recipient mice in which MHC II was specifically depleted on P2RY12^+^ microglia or TMEM119^+^ microglia (Fig. 7b and 7d). While EAU was successfully induced in control mice, in the absence of MHC II^+^ microglia, retinal inflammation was not amplified, and the peak EAU response was significantly reduced when compared to recipient mice expressing MHC II on microglia (Fig. 7b and 7d). Histological EAU evaluation of recipient mice with and without MHC II depletion on P2RY12^+^ or TMEM119^+^ cells at 13 days post adoptive transfer further confirmed that recipient mice with MHC II depletion on P2RY12^+^ or TMEM119^+^ cells failed to amplify retinal inflammation and achieve the peak EAU response when compared to recipient mice expressing MHC II on retinal microglia (Fig. 7c and 7e).

**Figure 7.**
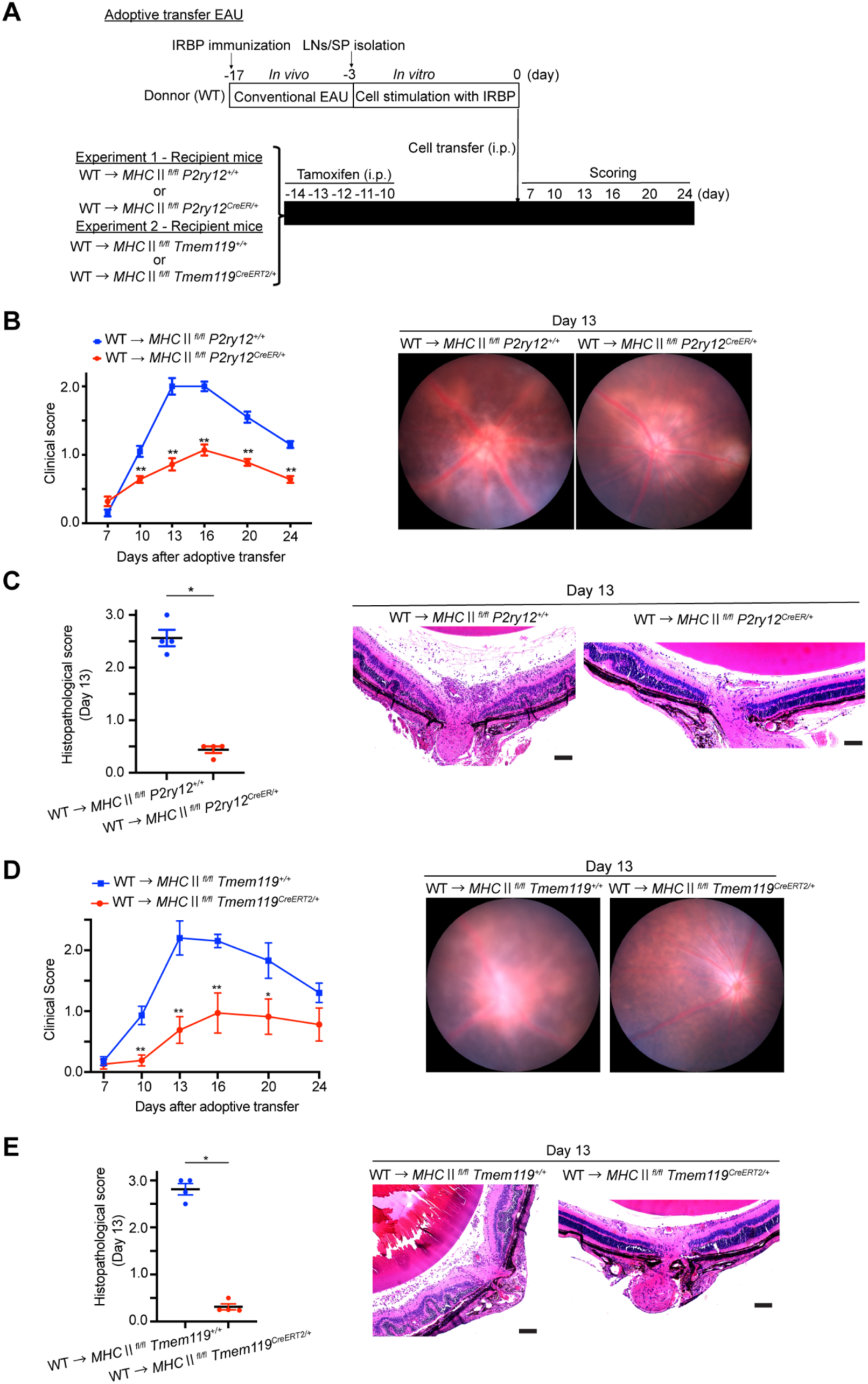
MHC II expression on microglia in recipient mice is required for the amplification of retinal inflammation in adoptive transfer EAU. (A) Schematic time course of adoptive transfer EAU experiment in which donor cells were isolated from WT mice at 14 days post immunization with IRBP, expanded in vitro, and administered to *MHC II^fl/fl^ P2ry12^+/+^* and *MHC II^fl/fl^ P2ry12^CreER/+^* mice or *MHC II^fl/fl^ Tmem119^+/+^*and *MHC II^fl/fl^ Tmem119^CreERT2/+^* and *MHC II^fl/fl^ CD11c^+/0^* mice. (B) The time course of EAU clinical scores in *MHC II^fl/fl^ P2ry12^+/+^* (n=5) and *MHC II^fl/fl^ P2ry12^CreER/+^* (n=7) recipient mice that received donor cells from WT and representative fundus images (day 13 post adoptive transfer). (C) Histopathological EAU score in *MHC II^fl/fl^ P2ry12^+/+^* (n=4) and *MHC II^fl/fl^ P2ry12^CreER/+^* (n=4) recipient mice on day 13 post adoptive transfer and representative histopathological images (H&E staining). Scale bar: 100 μm. (D) The time course of EAU clinical scores in *MHC II^fl/fl^ Tmem119^+/+^* (n=10) and *MHC II^fl/fl^ Tmem119^CreER/+^*(n=8) recipient mice that received donor cells from WT and representative fundus images (day 13 post adoptive transfer). (E) Histopathological EAU score in *MHC II^fl/fl^ Tmem119^+/+^* (n=4) and *MHC II^fl/fl^ Tmem119^CreER/+^* recipient mice on day 13 post adoptive transfer and representative histopathological images (H&E staining) (n=4 per group). Scale bar: 100 μm. All data are presented as mean ± SEM. All statistical analyses were performed using the Mann-Whitney U test. \**p*<0.05; \*\**p*<0.01.

Taken together, these results demonstrate that within the retina, MHC II^+^ microglia, but not infiltrating MHC II^+^ DCs, are required for the amplification and peak phases of EAU, indicating the local APC function of infiltrating DCs is insufficient for triggering the maximal response of retinal specific CD4^+^ T cells.

## Discussion

Microglia and retinal infiltrating DCs have two potential functions in the development of EAU (i) as APCs that present autoantigens via MHC II to infiltrating effector CD4^+^ T cells and (ii) as effector cells that directly and/or indirectly mediate destruction of retinal tissue through the release of inflammatory mediators and reactive oxygen species (ROS) and/or nitric oxide (NO) (32). This study focused on MHC II-restricted APC function of microglia and DCs within the retina during EAU and not on their effector functions. In our previous study, using the adoptive transfer model of EAU, we demonstrated that elimination of retinal microglia by treatment with PLX5622, an antagonist of CSF1R, prevented the development of EAU even though mice contained circulating IRBP-specific effector CD4^+^ T cells capable of mediating disease (21) (Fig. 8a). While these data clearly indicated microglia are essential for initiating uveitis, they were not informative about the antigen-presenting function of microglia. They did not reveal whether resident microglia and/or infiltrating DCs participated in presenting IRBP autoantigens to infiltrating effector CD4^+^ T cells during the different phases of EAU. Thus, the current study was designed to determine the role of antigen-presenting function, specifically in microglia and infiltrating DCs during EAU. This was achieved using *P2ry12*- and *Tmem119*-directed deletion of MHC II on microglia by a tamoxifen-inducible Cre-lox system and *CD11c*-directed deletion of MHC II on DCs.

**Figure 8.**
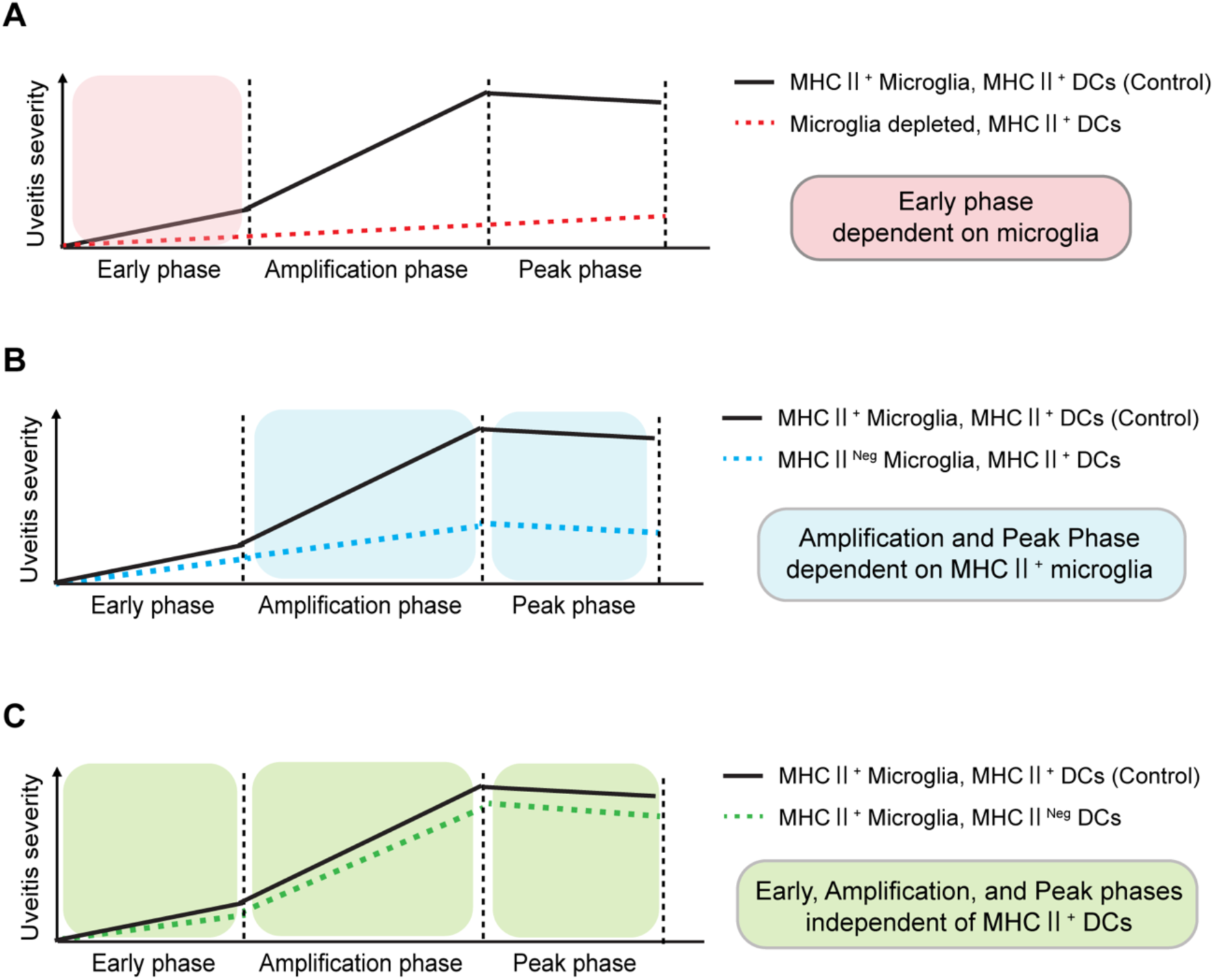
Graphical abstract. In the conventional and adoptive transfer models of EAU, the development of uveitis following the systemic induction (or adoptive transfer) of IRBP-specific T cells, can be divided into three phases: early, amplification, and peak. (A) Using the conventional model of EAU, we previously demonstrated that total microglia depletion prevents the early phase of uveitis, with no signs of uveitis detected, demonstrating the early phase of uveitis is dependent on microglia (20). (B) Specific depletion of MHC II on microglia has no effect on the early phase of uveitis when compared to control mice. However, the amplification and peak phases are significantly reduced, demonstrating the amplification and peak phases of uveitis are dependent upon MHC II^+^ microglia. (C) Using the adoptive transfer model of EAU, the specific depletion of MHC II on CD11c^+^ DCs has no effect on the development of uveitis, demonstrating all three phases of uveitis develop independently of MHC II^+^ DCs.

During EAU, MHC II^+^ cells were detected in the retina between day 10 and 20 post EAU induction and consisted of approximately equal numbers of microglia and DCs. The number of microglia dramatically increased during the amplification phase, presumably due to proliferation. Knockdown of MHC II, specifically on P2RY12^+^ / TMEM119^+^ microglia, prevented the peak phase of uveitis, and while a low level of retinal inflammation occurred, it was not amplified, and peak levels of retinitis were not achieved. We observed this in both the conventional and adoptive transfer methods of inducing EAU. These data indicate that without MHC II^+^ microglia, MHC II^+^ DCs are unable to mediate a peak inflammatory response in EAU (Fig. 8b). Finally, the severity of uveitis correlated with the number of MHC II^+^ microglia present within the retina. Together, these data indicate MHC II^+^ microglia are required for initiating and amplifying retinal inflammation during EAU.

It has long been thought that MHC II^+^ DCs are the primary APC in the spleen and lymph nodes that are responsible for the expansion of the antigen-specific CD4^+^ effector T cells that mediate EAU (9, 29), and previous studies showed inhibition of DC maturation, which prevents the upregulation of MHC II, attenuates EAU by preventing the generation of antigen-specific Th1 and Th17 cells (30, 31). However, our experiments are the first to genetically knockdown MHC II on DCs in EAU, confirming MHC II^+^ DCs as the APC responsible for the systemic induction and expansion of the antigen-specific CD4^+^ effector T cells in the conventional model of EAU. In mice with MHC II deficient DCs, it was not possible to induce IRBP-specific CD4^+^ effector T cells, and the mice failed to develop EAU.

It has been known for a long time that activated T cells freely infiltrate the retina, regardless of whether they recognize their cognate antigen, but if the activated T cells fail to recognize their cognate antigen, they are not reactivated, and their numbers quickly diminish without any evidence of inflammation (14). By contrast, if the activated T cells are triggered by their cognate antigen presented by APCs, then the T cells are reactivated, their numbers increase, and local inflammation is initiated. Therefore, a critical step that determines whether EAU develops is the initial presentation of autoantigens by APCs within the retina. However, determining whether these retinal APCs are resident microglia and/or infiltrating CD11c^+^ DCs has not been resolved because of the difficulty in identifying and separating microglia from infiltrating DCs. The solution to this problem emerged when Butovsky *et al.* (18) and Bennett *et al.* (19) reported the transcriptome study of homeostatic microglia in the brain, which identified specific markers (P2RY12 and TMEM119) that were enriched on microglia and allowed us to distinguish microglia from infiltrating DCs and other cells in the CNS. Using these microglia markers to drive the knockdown of MHC II specifically on microglia, we were able to determine their antigen presentation function during EAU.

Previous studies that indicated retinal microglia did <u>not</u> function as APCs during the development of EAU were based, in part, on the idea that microglia were important in maintaining the immune-privileged status of the retina (33). This idea was supported by *in vitro* studies of retinal microglia harvested from naive healthy mice, demonstrating that homeostatic microglia were unable to activate naïve T cells and could only weakly activate already primed T cells, as compared to splenic DCs and brain microglia (33). Activating retinal microglia *in vitro* by IFNγ treatment increased their ability to function as APCs but only slightly compared to similarly treated splenic DCs and brain microglia. Additional studies indicated that retinal microglia displayed immune inhibitory functions as shown by experiments in which homeostatic retinal microglia, when placed in co-cultures with splenic DCs, inhibited the ability of DCs to activate T cells. The inhibitory functions of retinal microglia were further supported by the observation that microglia express CD200R, a receptor for neuronal CD200 that triggers an anti-inflammatory signaling pathway (34). Together, these studies led to the idea that, in a normal homeostatic retina, microglia fail to function as APCs and possess immune inhibitory functions that help establish and maintain ocular immune privilege. Thus, it was thought that it would be difficult to overcome this inhibitory phenotype *in vivo* and convert microglia into MHC II^+^ APCs that could support the initiation and development of uveitis, implying that infiltrating DCs, with known potent antigen-presenting function, are the critical APCs in EAU.

By contrast, our results clearly demonstrate that microglia are the MHC II^+^ APCs that initiate and amplify retinal inflammation during EAU. Mice in which microglia were blocked from expressing MHC II did not achieve peak levels of retinitis, only a low level of retinal inflammation was observed, and this result was similar for mice in which either *P2ry12* or *Tmem119* was used to drive the knockdown of MHC II on microglia (Fig. 8b). The low level of inflammation observed in these mice with MHC II deficient microglia was most likely do to either a low level of residual MHC II on microglia because Cre recombination was less than 100% and/or the innate immune response mediated by MHC II^Neg^ microglia. These data, combined with our previous results indicating elimination of microglia by PLX5622 treatment completely blocked even the low level of inflammation observed (21) herein with MHC II^Neg^ microglia, indicate that retinal microglia function as APCs that trigger the early phase of EAU and, without the APC function of microglia, no retinitis will develop, even in the presence of circulating IRBP-specific effector T cells.

EAU and experimental autoimmune encephalitis (EAE) are CNS autoimmune diseases that have been frequently compared for similarities in pathogenesis. This has been informative for certain situations (35, 36), even though there are obvious and significant differences between retinal and brain anatomical structure and resident cell populations. However, regarding the antigen-presenting function of MHC II^+^ APCs, EAE is not helpful because whether brain microglia function as MHC II^+^ APCs is still unresolved, with some studies indicating they function as APCs (37) and others indicating they do not (38).

Our somewhat surprising result was that retinal infiltrating MHC II^+^ DCs do not function as APCs at any phase of EAU development, as shown by the fact that *CD11c*-driven knockdown of MHC II on retinal DCs had no negative impact on the amplification and/or peak phases of EAU (Fig. 8c). The idea that infiltrating DCs served as APCs in EAU was first proposed in historical papers that indicated retinal infiltrating DCs were the primary APC during EAU, using the technology available at the time to deplete infiltrating DCs, such as dichlorodimethylene diphosphonate containing liposomes, which lacked a high degree of cell specificity (39). However, more recent studies by McPherson *et al.* provided supporting evidence that DCs have an important role as APCs in EAU. Using a spontaneous autoimmune uveitis model system that develops in R161H mice (on the B10.RIII background), they crossed these mice to *CD11c^DTR/GFP^* mice (on the B6/J background) to produce F1 mice (*R161H^+/-^* x *CD11c^DTR/GFP^*) (40). These mice were then used in a parabiosis model system that connected the vasculature of the F1 mice with WT recipient mice (B10.RIII × B6/J F1 mice). This approach allowed them to track retinal infiltration of GFP^+^ CD11c^+^ DCs during the development of uveitis in the WT recipient mice. Studying 29 pairings, GFP^+^ cells were detected in 19 recipient mice, of which 12 developed uveitis. However, uveitis never developed in the 10 mice that lacked infiltrating GFP^+^ cells. The authors concluded that the recruitment of APCs from the circulation was a prerequisite for the induction of spontaneous uveitis. While this is strong circumstantial evidence that recruitment of circulating APCs is required for the development of uveitis, these experiments did not directly test the antigen-presenting function of the recruited APCs. Furthermore, the recruited GFP^+^ cells may have an MHC II-independent role in mediating tissue destruction through ROS and/or NO production. This scenario would be consistent with our results, where we observed in mice with MHC II deficient CD11c^+^ cells, that uveitis developed normally and was not different from WT controls. Since we did not block infiltration of the MHC II deficient CD11c^+^ cells in our experiments, these cells may still have participated in tissue destruction independent of antigen presentation. However, the experiments by McPherson *et al.* (40) were not consistent with our previous results that no EAU developed in the absence of microglia by PLX5622 treatment, indicating that microglia controlled the initial signals that trigger the start of uveitis, and the disease process could not be initiated by infiltrating DCs alone. However, it is difficult to directly compare our results with the parabiosis model since the latter uses mice that spontaneously develop uveitis, and our model requires immunization of IRBP antigens along with adjuvants that trigger potent innate immunity. Therefore, these two models may differ in the initial phases of disease induction.

While our results identified the cells that serve as MHC II^+^ APCs during the development of EAU, there are still many unanswered questions raised by our results. Microglia are known as the gatekeepers of the retinal microenvironment and contribute to maintaining immune privilege. How do homeostatic microglia transition from MHC II^Neg^ CD200R^+^ inhibitory cells, with little or no ability to activate T cells, to become activated MHC II^+^ APCs capable of presenting autoantigens and reactivating infiltrating T cells to induce escalating levels of retinal inflammation? Why do MHC II^+^ DCs fail to participate in the presentation of autoantigens during the different phases of disease? Recent data using lineage tracing indicates that retinal microglia and infiltrating monocytes are heterogeneous (41), which raised the question that these subpopulations have equal antigen-presenting functions during the development of EAU.

In conclusion, our results indicate that microglia are essential during all three phases of EAU, in the early phase when microglia transition into MHC II^+^ APCs and in the amplification and peak phases. Unexpectedly, while retinal infiltrating MHC II^+^ CD11c^+^ DCs were present within the retina, their antigen-presenting function was not required for the early, amplification, and peak phases of EAU. This indicates that microglia are an important potential therapeutic target that can prevent and/or diminish uveitis even in the presence of circulating IRBP-specific CD4^+^ effector T cells.

## Materials and Methods

### Mice

All animal experiments adhered to the guidelines of the ARVO Statement for the Use of Animals in Ophthalmic and Vision Research and were approved by the Animal Care Committee of the Massachusetts Eye and Ear Infirmary. *MHC II^fl/fl^*mice (Strain #:013181), *CD11c-Cre* mice (Strain #:008068), *Tmem119-CreERT2* mice (Strain #:031820), and *P2ry12-CreER* mice (Strain #:034727), were obtained from the Jackson Laboratory (Bal Harbor, ME, USA). To generate *MHC II^fl/fl^CD11c-Cre*, *MHCⅡ^fl/fl^ Tmem119-CreERT2*, and *MHC II^fl/fl^ P2ry12-CreER* mice, *MHC II^fl/fl^* mice were crossed with *CD11c-Cre*, *Tmem119-CreERT2,* and *P2ry12-CreER* mice, respectively. Mice were fed standard laboratory chow and allowed free access to food and water in a climate-controlled room with a 12-hour light/12-hour dark cycle. All mice used for the experiments were 7-10 weeks old. For anesthesia, tribromoethanol solution (12.5 mg/ml) was injected intraperitoneally for survival procedures (250 mg/kg) and non-survival procedures (400 mg/kg).

### Induction of EAU

For the induction of conventional EAU, 200 μg of human IRBP-p (peptide 1-20) (Biomatik, Wilmington, DE) was emulsified in Complete Freund’s Adjuvant (1:1 w/v) (Difco, Detroit, MI) containing *M. tuberculosis,* H37Ra (Difco, Detroit, MI) (5.0 mg/ml). A total of 200 μL emulsion was injected subcutaneously into the neck (100 μL), top of the right foot (50 μL), and left inguinal region (50 μL). Concurrent with immunization, 100μL of pertussis toxin solution (Sigma-Aldrich, St. Louis, MO) (10 μg/ml) was injected intraperitoneally. For analgesia, 150 μL of buprenorphine hydrochloride solution (Reckitt Benckiser, Slough, United Kingdom) (15μg/ml) was injected subcutaneously into the neck immediately and 3 hours after emulsion administration.

Adoptive transfer of EAU was induced as previously described (20), with minor modifications. Briefly, donor mice were immunized as described above, and at 14 days after immunization, the spleens (SP) draining lymph nodes (LN) (cervical, axillary, and inguinal) were collected. Lymphocytes from SP and draining LNs were cultured in the presence of IRBP-p (10 μg/ml) and IL-23 (R&D systems, Minneapolis, MN) (10 ng/ml) for 72 hours in RPMI 1640 (Thermo Fisher Scientific, Waltham, MA) supplemented with FBS (Thermo Fisher Scientific, Waltham, MA) (10 %), HEPES (Thermo Fisher Scientific, Waltham, MA) (10 mM), MEM Non-Essential Amino Acids Solution (Thermo Fisher Scientific, Waltham, MA) (1×), Sodium Pyruvate (Thermo Fisher Scientific, Waltham, MA) (1 mM), 2-Mercaptoethanol (Sigma-Aldrich, St. Louis, MO) (50 μM), and penicillin-streptomycin (Thermo Fisher Scientific, Waltham, MA) (100 U/ml). The non-adherent cells in suspension were transferred to new dishes on days one and two of culture. On day three, activated lymphocytes were purified by gradient centrifugation using Histopaque-1083 (Sigma-Aldrich, St. Louis, MO, USA). Finally, 200 μL of the cell suspension (5×10^6^ cells/mouse) was injected intraperitoneally into recipient mice.

### Assessment of EAU

Fundus images were taken for clinical assessment using a MICRON IV (Phoenix-Micron, Bend, OR, USA). The clinical score of each mouse was graded in a blinded manner on a scale of 0-4 in half-point increments, as described previously (42). Trace chorioretinal lesions and/or minimal vasculitis were scored as 0.5. Mild vasculitis and/or small focal chorioretinal lesions (≤ 5) were scored as 1. Severe vasculitis and/or multiple chorioretinal lesions (>5) were scored as 2. Linear chorioretinal lesions, subretinal neovascularization, and hemorrhages were scored as 3. Retinal inflammation with a large retinal detachment and severe hemorrhages were scored as 4. For histological assessment, enucleated eye was fixed in a 10% neutral buffered formalin solution (Sigma-Aldrich, St. Louis, MO) overnight. The fixed tissues were embedded in paraffin, thinly sliced (5μm), deparaffinized, and stained with H&E. The severity of EAU was scored in a blinded manner on a scale of 0-4 in half-point increments, as described previously (43). Briefly, focal non-granulomatous monocytic infiltration in the choroid, ciliary body, and retina was scored as 0.5. Monocytic infiltration in the vitreous and retinal perivascular was scored as 1. Granuloma formation in the uvea and retina, occluded retinal vasculitis, photoreceptor folds, serous retinal detachment, and loss of photoreceptors was scored as 2. Granulomatosis formation at the level of the retinal pigment epithelium and the development of subretinal neovascularization were scored as 3 and 4, respectively, according to the number and size of the lesions (44). The average of scores from both eyes was determined as the score for the animal.

### Microglia-specific depletion of MHC II

MHCII depletion in microglia was accomplished using *MHC II^fl/fl^ Tmem119-CreERT2* mice whose Cre-ERT2 fusion gene was driven by *Tmem119* promoter, and *MHC II^fl/fl^*P2RY12*-CreER* mice whose Cre-ER fusion gene was driven by P2RY12 promoter. In these transgenic mice, Cre recombinase activation was induced by tamoxifen under the control of a specific promoter (*P2RY12* or *Tmem119*). Tamoxifen (Sigma-Aldrich, St. Louis, MO) was dissolved in corn oil at 20 mg/ml. At 14 days prior to EAU induction, 100 µL of the tamoxifen solution was injected intraperitoneally for five consecutive days.

### Delayed-type Hypersensitivity Measurement

Antigen-specific delayed hypersensitivity was assessed as previously described (45). On day 19 after immunization, mice were intradermally injected with 10 μL of IRBP-p suspended in PBS (1.0 mg/ml) into the pinna of the right ear. Ear swelling was measured after 48 hours using a micrometer (Mitutoyo, Kanagawa, Japan). Delayed hypersensitivity was calculated as the difference in ear thickness before and after the challenge. The following formula was used for calculation: [(48-hour measurement – 0-hour measurement) in the test ear] - [(48-hour measurement – 0-hour measurement) in the control ear].

### Cell Proliferation Assay

At 14 days post EAU induction, the SP and draining LNs were collected, and cells were suspended at 2×10^5^ cells/200μL of medium in 96-well flat-bottom plates. The cells were cultured in triplicate in the presence of IRBP-p (10 μg/ml), Con A (Sigma-Aldrich, St. Louis, MO) (1.0 μg/ml), or culture medium only. During the last 4 hours of the 72-hour culture, 100 μL of culture supernatant was removed, 10μL of Cell Counting Kit-8 (DOJINDO, Kumamoto, Japan) was added to each well, and cells were cultured for an additional four hours. At the end of the culture period, the number of viable cells in each well was measured as the absorbance (450 nm) of reduced WST-8 (2-(2-methoxy-4-nitrophenyl)-3-(4-nitrophenyl)-5-(2,4-disulfophenyl)-2H-tetrazolium, monosodium salt) using a Spectra Max M3 Multi-Mode Microplate Reader (Molecular Devices, San Jose, CA).

### Flow Cytometric Analysis of lymph nodes, spleen, and retina

For LN and SP single cell suspensions, the draining LNs and SP were collected and placed in a 70μm cell strainer and homogenized. The cells were then suspended in RBC lysis buffer (BioLegend, San Diego, CA, USA) (1×) for 2 min at room temperature and the cell pellet was suspended in 1×PBS supplemented with BSA (Sigma-Aldrich, St. Louis, MO) (0.5%) and UltraPure 0.5 M EDTA, pH 8.0 (Thermo Fisher Scientific, Waltham, MA) (2mM). For retinal single cell suspensions, retina was collected and incubated for 10 minutes at 37 ℃ in DPBS (Thermo Fisher Scientific, Waltham, MA) supplemented with L-cysteine (Sigma-Aldrich, St. Louis, MO) (100 mg/ml), sodium hydroxide (Sigma-Aldrich, St. Louis, MO) (1.3 mM), and papain (Worthington Biochemical, Lakewood, NJ) (10 U/ml). Following the incubation with a mixed solution of ovomucoid (Worthington Biochemical, Lakewood, NJ) (1.5 mg/ml) and DNase (Worthington Biochemical, Lakewood, NJ) (0.1 mg/ml) for 3 minutes at room temperature to stop the enzymatic digestion, retina was homogenized by gentle pipetting and transferred to a 50ml tube through a 70μm cell strainer. The cell pellet was suspended in 1×PBS supplemented with BSA (Sigma-Aldrich, St. Louis, MO) (0.5 %) and UltraPure 0.5M EDTA, pH 8.0 (Thermo Fisher Scientific, Waltham, MA) (2 mM).

Single cell suspensions were blocked with anti-mouse CD16/32 mAb (BioLegend, San Diego, CA) for 15 minutes at 4℃. Dead cells were stained with 4’,6-diamidino-2-phenylindole (DAPI) (Thermo Fisher Scientific, Waltham, MA) (1.0 μg/ml) or a LIVE/DEAD fixable dead stain kit (violet) (Thermo Fisher Scientific, Waltham, MA). The following antibodies were used for staining: Alexa Fluor 488 anti-mouse CD3 (17A2) (1:100) (BioLegend, San Diego, CA), PE anti-mouse CD4 (GK1.5) (1:100) (BioLegend, San Diego, CA), BB515 anti-mouse CD11b (M1/70) (1:100) (BD, Ashland, OR), BB515 anti-mouse CD11c (N418) (1:100) (BD, Ashland, OR), Brilliant Violet 785 anti-mouse CD45 (30-F11) (1:50) (BioLegend, San Diego, CA), PE anti-mouse I-A/I-E (M5/114.15.2) (1:100) (BioLegend, San Diego, CA), and APC anti-mouse P2RY12 (S16007D) (1:100) (BioLegend, San Diego, CA). For TMEM119 staining, rabbit anti-mouse Tmem119 (106–6) (1:500) (Abcam, Waltham, MA) and APC-conjugated goat anti-rabbit IgG (H+L) antibody (1:500) (Thermo Fisher Scientific, Waltham, MA, USA) was used as the primary and secondary antibodies, respectively. Single color staining and fluorescence minus one sample were used for gain setting, gating, and compensation. Acquired data were analyzed using FlowJo software 10.8.1 (BD, Ashland, OR).

### Mixed lymphocyte proliferation assay

Draining LNs and SP were isolated from conventional EAU-induced *MHC II^fl/fl^ CD11c^Cre/0^* and *MHC II^fl/fl^ CD11c^+/0^* mice 14 days after EAU induction. CD4^+^ T cells was isolated from LNs using a CD4^+^ T cell isolation kit (Milteyni Biotech, Bergisch Gladbach, Germany) (Fig. S4A, B, and C). CD11c^+^ DCs was isolated from SP using CD11c MicroBeads UltraPure (Milteyni Biotech, Bergisch Gladbach, Germany) (Fig. S4D, E, and F). The purified CD4^+^ T cells were stained with Far-Red using CellTrace Cell Proliferation Kit (Thermo Fisher Scientific, Waltham, MA) before coculture with CD11c^+^ DCs. Aliquots of the Far-Red-stained CD4^+^ T cells (2×10^5^ cells) were co-cultured with CD11c^+^ DCs (1×10^5^ cells) in U-bottom 96-well plates (200 μl/well) in the presence of IRBP-p (10 μg/ml) for three days. After three days of co-culture, the frequency and number of proliferating CD4^+^ T cells in each well was determined by diluted Far-Red expression gated on live CD4^+^ cells. The percentage and total number of proliferating CD4^+^ T cells was determined in each coculture combination.

### Statistical Analysis

Data are presented as mean ± standard error of the mean (SEM). The difference between the two groups was analyzed using the Mann-Whitney U test. Multiple-group comparison was analyzed using a one-way analysis of variance with Tukey’s multiple comparison test (\**p*<0.05, \*\**p*<0.01, \*\*\*\**p*<0.001, and \*\*\*\**p*<0.0001 throughout the paper). Spearman’s rank correlation coefficient was calculated to examine the relationship between two variables. All statistical analyses were performed using Prism10 software (GraphPad Software, San Diego, CA, USA).

## Acknowledgments

We thank all our laboratory members for their fruitful discussions and for their help with the experiments. We also thank the technical staff at the Schepens Eye Research Institute of Massachusetts Eye and Ear, Harvard Medical School, for supporting this research. All Figures were created with Adobe Illustrator (Adobe, San Jose, CA, USA). This study was supported by the National Institute of Health/National Eye Institute: Grant R01EY031291 (M.G.K. and B.R.K.) and Core Grant (P30EY003790).

## Supplementary Figures

**Fig. S1.**
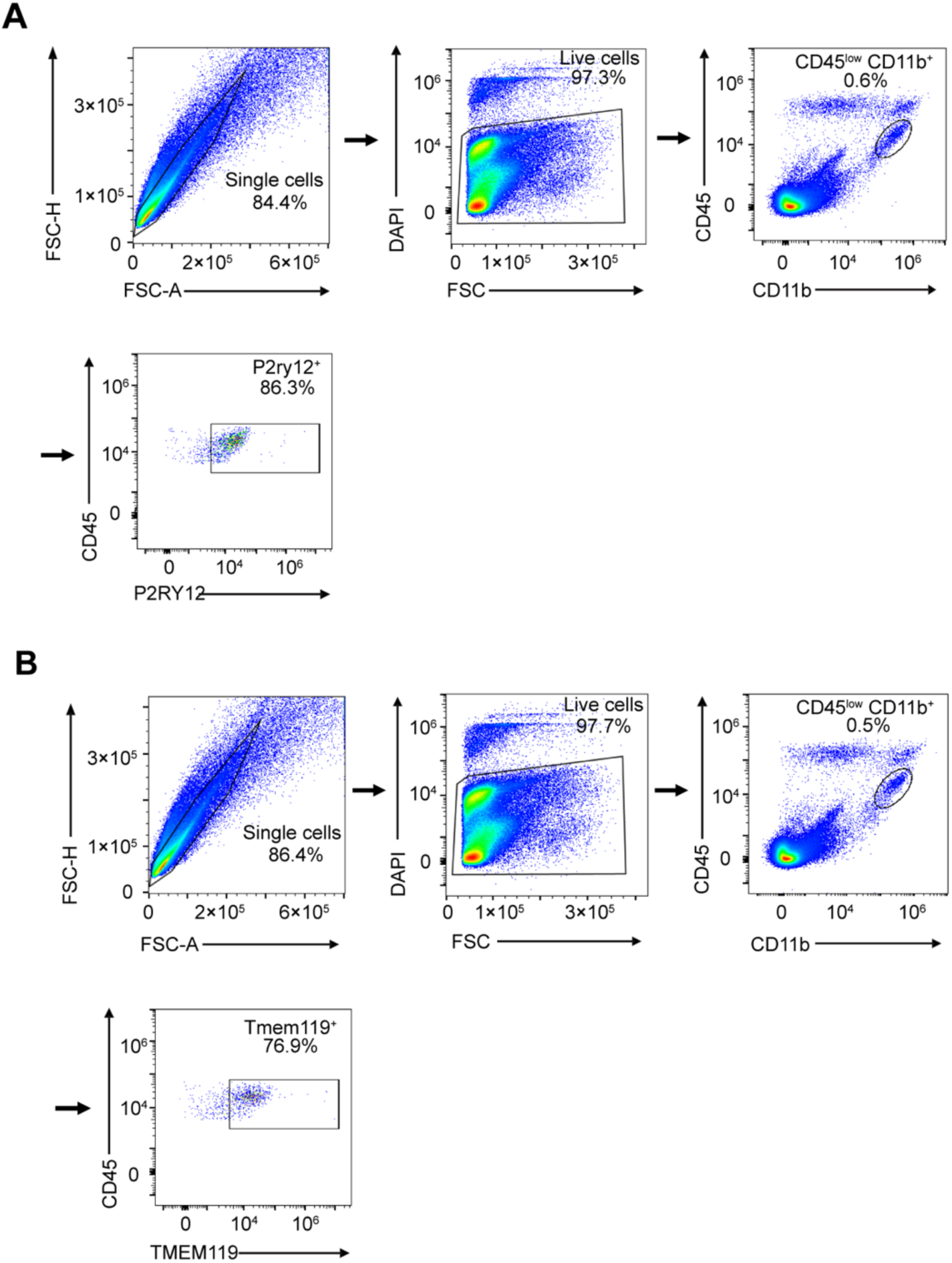
Complete gating strategy for flow cytometric analysis of retinal microglia. (A and B) Representative flow cytometric analysis of retinal microglia in naïve C57BL/6J WT mice. Following the preparation of a single-cell suspension, FSC-A vs. FSC-H was used to gate on singlets and exclude doublets. Within the singlet gate, DAPI staining was used to exclude dead cells. Within the live cell gate, microglia were identified as CD45^low^ CD11b^+^ and further defined by (A) P2RY12 expression and (B) TMEM119 expression.

**Fig. S2.**
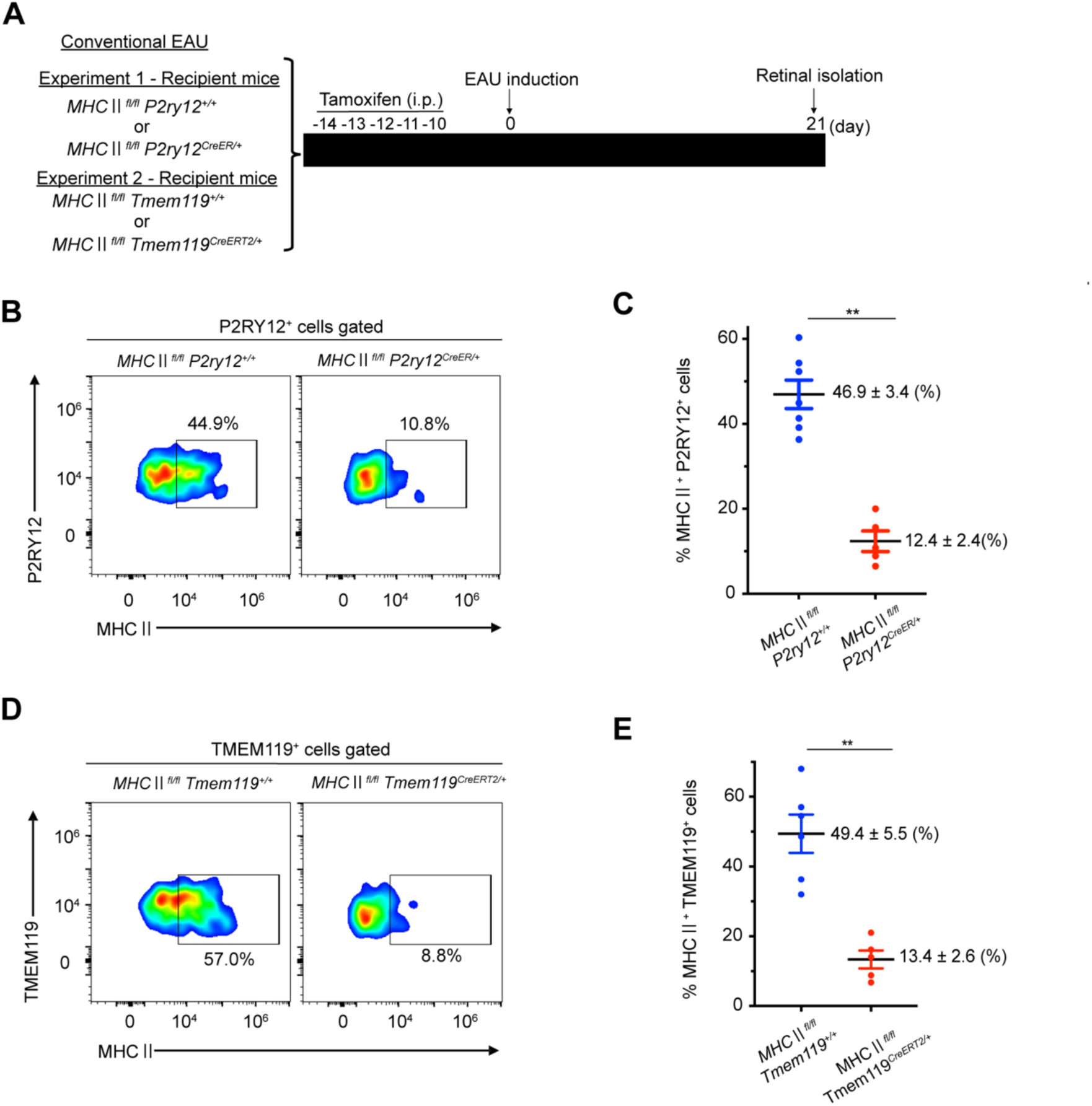
Confirmation of MHC II knockdown in retinal microglia from *MHC II^fl/fl^ P2ry12^CreER/+^*and *MHC II^fl/fl^ Tmem119^CreERT2/+^* mice. (A) Schematic time course of conventional EAU experiment in which Tamoxifen administration was initiated 14 days before immunization with IRBP to specifically knock out MHC II on microglia, using *MHC II^fl/fl^ P2ry12^CreER/+^* and *MHC II^fl/fl^ Tmem119^CreER/+^* mice. At 21 days post IRBP immunization, retinas were collected and analyzed by flow cytometry. (B) Representative flow cytometry plot showing MHCⅡ expression on P2RY12^+^ cells in the retina collected from *MHC II^fl/fl^ P2ry12^+/+^* (n=7) and *MHC II^fl/fl^ P2ry12^CreER/+^* (n=5) mice 21 days after EAU induction. (C) Quantitative analysis of MHC II^+^ P2RY12^+^ cells. (D) Representative flow cytometry plot showing MHC II expression on TMEM119^+^ cells in the retina collected from *MHC II^fl/fl^ Tmem119^+/+^* (n=6) and *MHC II^fl/fl^ Tmem119^CreER/+^* (n=5) mice 21 days after EAU induction. (E) Quantitative analysis of MHC II^+^ TMEM119^+^ cells. All data are presented as mean ± SEM. All statistical analyses were performed using the Mann-Whitney U test. \*\**p*<0.01.

**Fig. S3.**
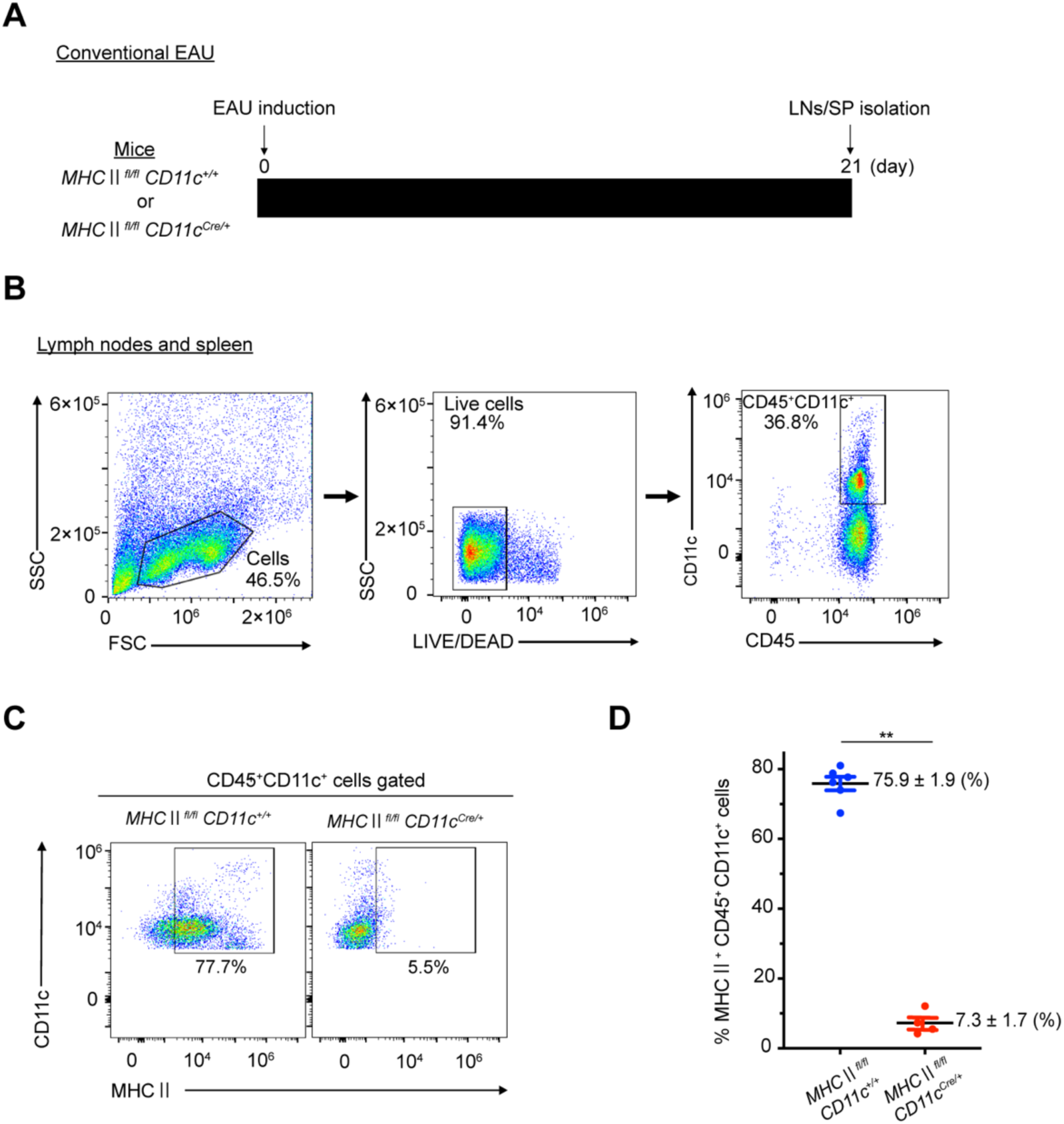
Gating strategy for flow cytometric analysis of CD11c^+^ dendritic cells and confirmation of MHC II knockdown in CD11c^+^ dendritic cells from *MHC II^fl/fl^ CD11c^+/0^*and *MHC II^fl/fl^ CD11c^Cre/0^* mice. (A) Schematic time course of conventional EAU experiment in which *MHC II^fl/fl^ CD11c^+/0^* or *MHC II^fl/fl^ CD11c^Cre/0^* mice were immunized with IRBP. At 21 days post IRBP immunization, lymph nodes (LNs) and spleen (SP) were collected and analyzed by flow cytometry. (B) Following preparation of a single-cell suspension, FSC vs SSC was used to exclude debris. LIVE/DEAD stain was used to DAPI exclude dead cells. Within the live cell gate, CD11c^+^ dendritic cells were identified as CD45^high^ CD11c^+^. (C) Representative flow cytometry plot showing MHC II expression CD45^+^ CD11c^+^ cells in the LN/SP single cell suspension from *MHC II^fl/fl^ CD11c^+/0^* (n=6) and *MHC II^fl/fl^ CD11c^Cre/0^* (n=4) mice. (D) Quantitative analysis of MHC II^+^ expression on CD45^+^ CD11c^+^ cells. All data are presented as the mean ± SEM. All statistical analyses were performed using the Mann-Whitney U test. \*\**p*<0.01.

**Fig. S4.**
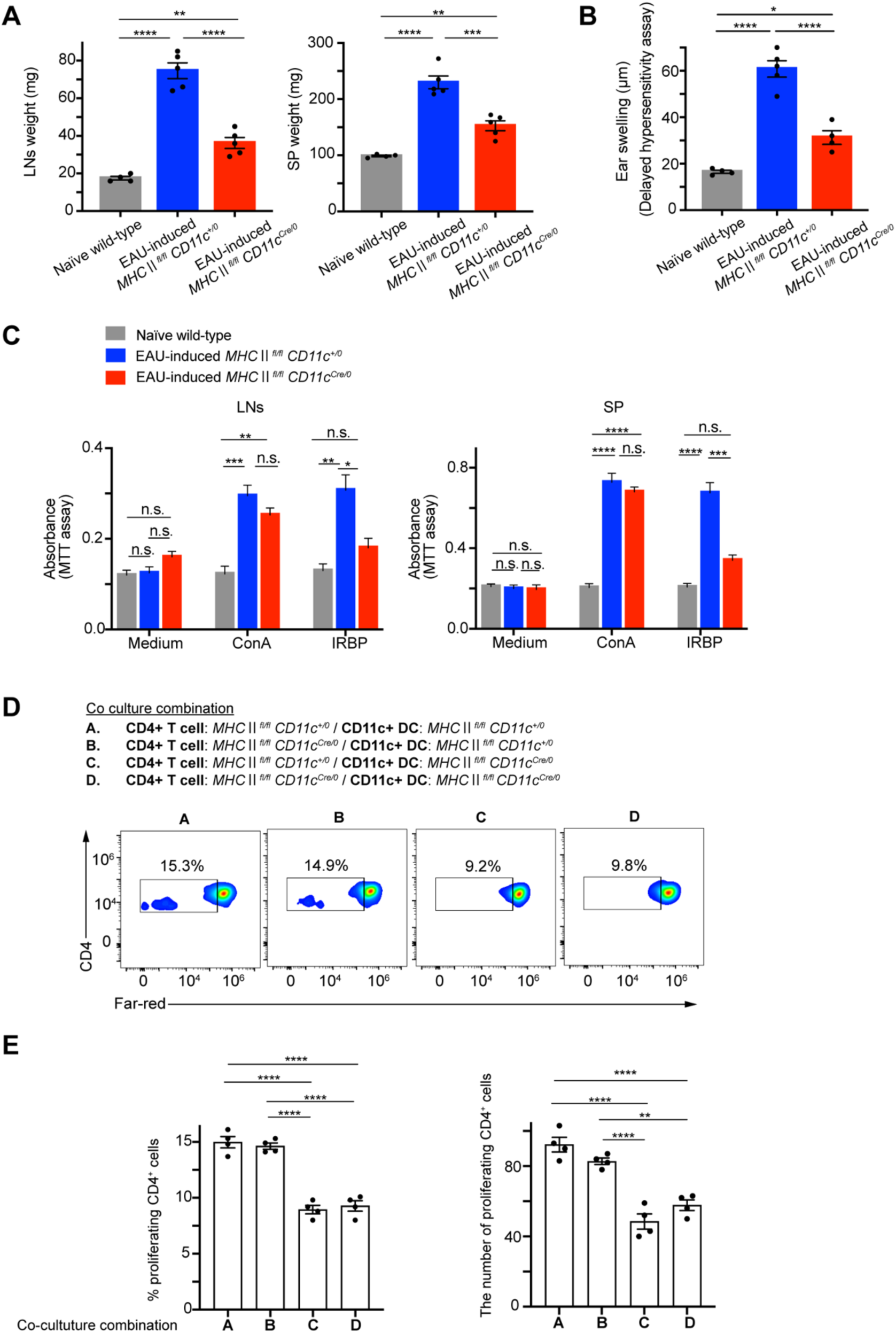
CD11c^+^ dendritic cell induces an antigen-specific systemic immune response by leading to the proliferation of CD4^+^ T cells. (A) Weight of draining lymph nodes (LNs) and spleen (SP) from *MHC II^fl/fl^ CD11c^+/0^* mice (n=5) and *MHC II^fl/fl^ CD11c^Cre/0^* (n=5) mice on day 21 post-IRBP immunization. Naïve WT mice (n=4) served as controls. (B) Antigen-specific delayed-type hypersensitivity, as determined by ear swelling, was evaluated in naïve wild-type mice (n=4) and *MHC II^fl/fl^ CD11c^+/0^* mice (n=5) and *MHC II^fl/fl^ CD11c^Cre/0^* (n=5) mice at day 21 post-IRBP immunization. Mice were injected intradermally with IRBP into the pinna of one ear on day 19, and ear swelling was measured after 48 h using a micrometer. (C) Antigen-specific (IRBP) and non-specific (ConA) cell proliferation was evaluated by MTT assay using cells isolated from LNs and SP of *MHC II^fl/fl^ CD11c^+/0^* mice (n=4) and *MHC II^fl/fl^ CD11c^Cre/0^* (n=4) mice on day 21 post-IRBP immunization. Naïve WT mice (n=4) served as controls. (D) T cell proliferation was also examined in co-cultures containing purified CD4^+^ T cells (stained with Far-Red) and CD11c^+^ DCs isolated from *MHC II^fl/fl^ CD11c^+/0^* (n=4) and *MHC II^fl/fl^ CD11c^Cre/+^* (n=4) mice on day 21 post IRBP immunization. The T cells and DCs were cultured in a medium containing the IRBP for three days using four different co-culture combinations. After three days of co-culture, CD3^+^ CD4^+^ cells with reduced Far-Red expression were defined as proliferating CD4^+^ T cell. Representative flow cytometry plots showing the percentage of Far-red^+^ CD4^+^ T cells in each co-culture combination. (E) Quantitative analysis of the percentage and total number of proliferating CD4^+^ T cells in each coculture combination. All data are presented as mean ± SEM. All statistical analyses were performed using a one-way analysis of variance with Tukey’s multiple comparison test. \**p*<0.05; \*\**p*<0.01; \*\*\*\**p*<0.001; \*\*\*\**p*<0.0001.

**Fig. S5.**
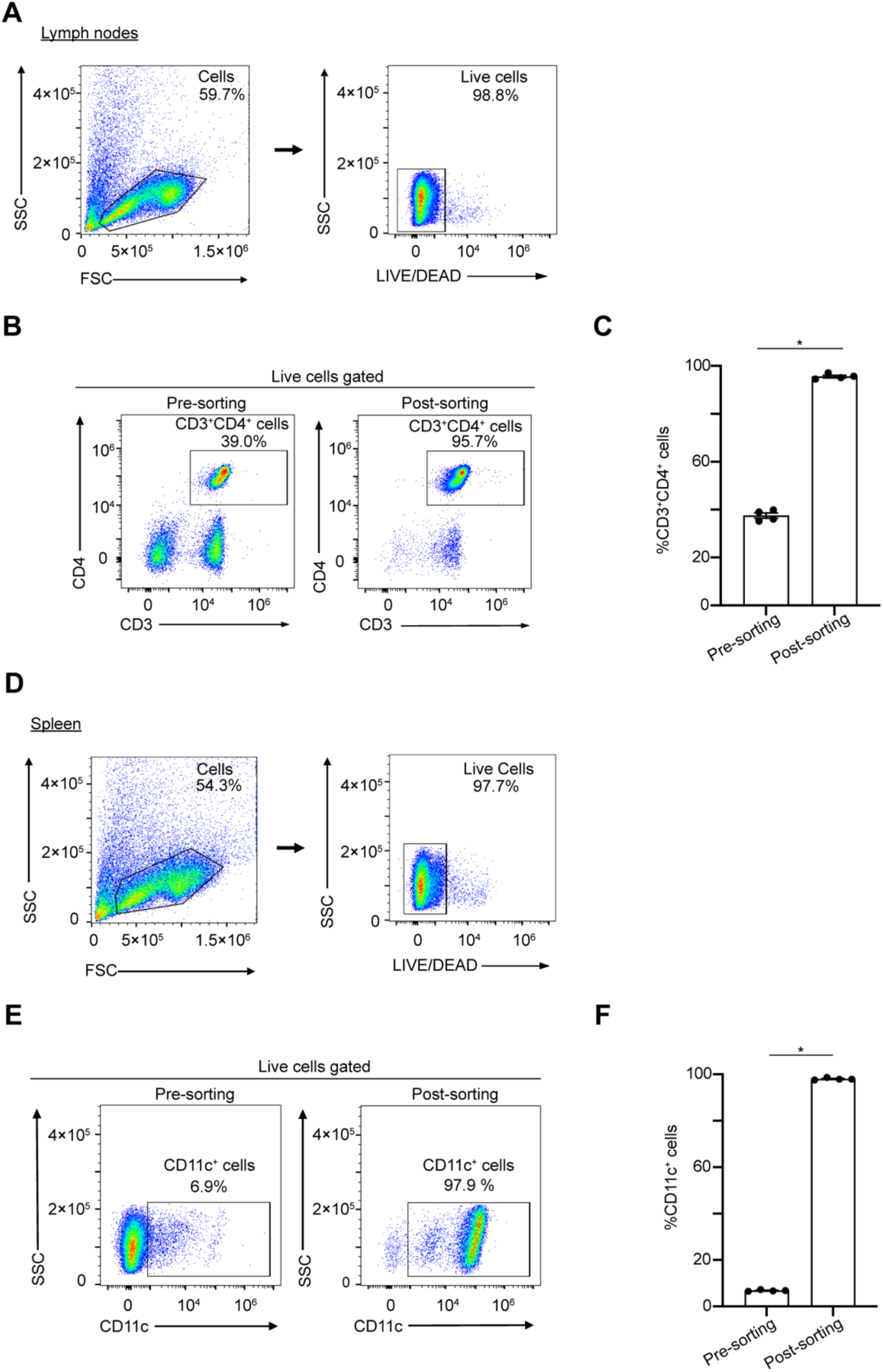
Purification of CD4^+^ T cells and CD11c^+^ cells using a magnetic-activated cell sorting system. (A) Gating strategy to identify live cells in the lymph nodes (LNs) of naïve wild-type mice. (B) Representative flow cytometry plots showing CD4 and CD3 expression in the LN. (C) The percentage of CD3^+^ CD4^+^ cells in the LNs of naïve wild-type mice, pre-sort (n=4) and post-sort (n=4). (D) Gating strategy to identify live cells in the spleen (SP) of naïve wild-type mice. (E) Representative flow cytometry plots showing CD11c expression in the SP. (F) The percentage of CD11c^+^ cells in the SPs of naïve wild-type mice, pre-sort (n=4) and post-sort (n=4). All data are presented as mean ± SEM. All statistical analyses were performed using the Mann-Whitney U test. \**p*<0.05.

